# Baseline soil carbon, time since conversion and climate, not system characteristics, shape soil carbon responses in Neotropical agroforestry. A meta-analysis

**DOI:** 10.64898/2026.02.06.704434

**Authors:** Damien Beillouin, Cloé Verstraete, Rémi Cardinael, Ulysse Chabroux, Jean-Baptiste Laurent, Pierre-André Waite, Julien Demenois

## Abstract

Agroforestry is promoted as a nature-based climate solution. Yet, the size of soil organic carbon gains and the factors controlling them remain uncertain, particularly in the tropics, where much of the evidence lies in grey and non-English literature. Here we analysed 322 response ratios and 277 storage rates from 54 Neotropical field studies with three-level meta-analyses, meta-regressions and gradient-boosted trees. Agroforestry increased soil organic carbon stocks by 6.8% (95% CI: 1.7−12.2%), or 0.25 (0.05−0.45) Mg C ha^-^¹ yr^-^¹. The large heterogeneity in the response was mostly explained by biophysical conditions (16 to 23%), whereas agroforestry type, previous land use, number of tree species and crop contributed little. Relative gains declined with increasing initial soil carbon, from 19% at 20 Mg C ha^-^¹ to 1.3% at 90 Mg C ha^-^¹, and storage rates followed the same pattern. The absolute amount of carbon added, 2.9 Mg C ha^-^¹ (0.3−5.5), did not vary with initial stock. The decline in relative gains was more pronounced at warmer sites. Time since conversion was the second strongest driver: relative gains increased by 0.4% per year, whereas annual storage rates fell from 1.2 Mg C ha^-^¹ yr^-^¹ at 2 years to 0.1 at 10. Results were the same in English and Spanish publications, and in peer-reviewed and grey literature, and no single study drove the relative gains. Only seven studies went beyond 20 years, and deeper layers were sampled only in young systems. Here we show for the first time that the relative soil carbon response of Neotropical agroforestry is governed by the initial state of the soil and by climate, not by the system typologies used in carbon accounting. Targeting carbon-depleted soils maximises the relative gain, not the amount stored, which increased with system age.

## 1 Introduction

Agroforestry is included in climate pledges and global mitigation assessments as a way to increase soil carbon (Bossio et al. 2020; Roe et al. 2021; Fig. 1). Yet the estimates behind these assessments vary widely, from 0.08 to 5.7 Gt CO^-^ -eq yr^-^¹ (Griscom et al. 2020). Field measurements are similarly variable, ranging from net losses of 1.15 to gains of 0.75 Mg C ha^-^¹ yr^-^¹ (Cardinael et al. 2018), with particularly large variation among tropical systems (Nair et al. 2009; Becker et al. 2025).

**Fig. 1.**
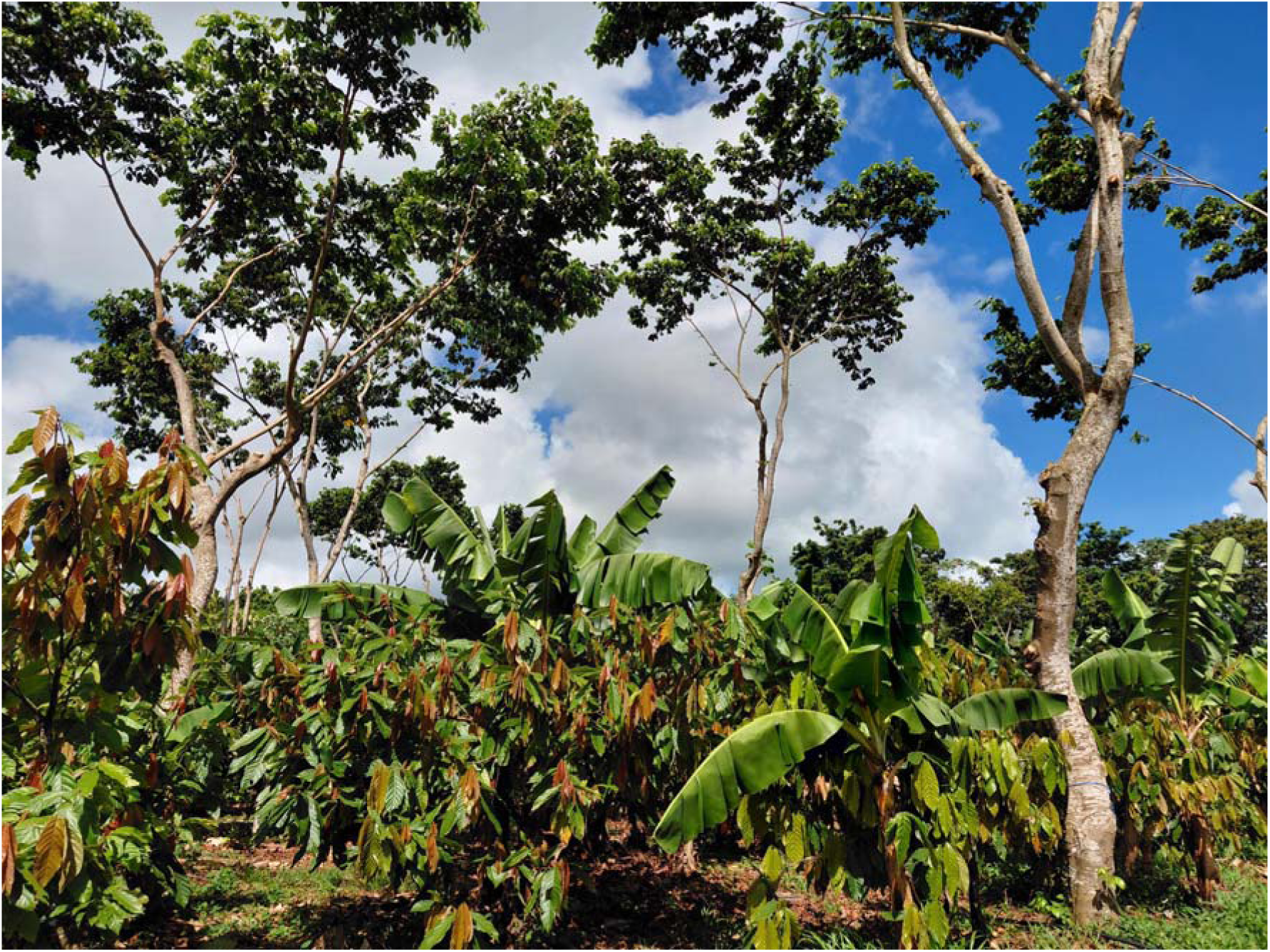
Agroforestry system combining cocoa (*Theobroma cacao* L.), banana (Musa sp.), and *Inga ingoides* (Rich.) Willd. at the Banabio experiment, Martinique**. Photocredit: C.-M. Rohé (Cirad).**

Responses of soil organic carbon (SOC) are often non-linear, jointly governed by baseline soil carbon stocks, land-use legacies and actual carbon inputs, which set both the rate and the persistence of accumulation (Basile-Doelsch et al. 2020; Terrer et al. 2021). Yet, studies are often short-term and focused on topsoil, and much of the literature is scattered across grey and non-English sources (Amano et al. 2023). Moreover, synthesis approaches usually describe agroforestry systems through broad categories, making it difficult to capture variation among sites and to assess whether relationships found in the available studies hold elsewhere (Terasaki Hart et al. 2023). In addition, the drivers of carbon change themselves covary further complicating precise estimation of agroforestry effects on soil organic carbon especially in sparse, unbalanced datasets (Altman and Royston 2006; Powers et al. 2011; Sun et al. 2023). As a result, scaling the available evidence to robust and durable estimates of global mitigation potential remains challenging (Friedlingstein et al. 2019; Cook-Patton et al. 2021).

This challenge is particularly acute across the Neotropics, where strong climatic, edaphic, and land-use gradients amplify SOC variability (Guevara et al. 2018), and where the geographic bias of field observations limits spatial extrapolation (Powers et al. 2011). Moreover, over the past two decades, a 50% increase in extreme weather events across Latin America and the Caribbean (LAC) has destabilized terrestrial carbon pools, increasing the risk of soil carbon reversal and undermining the permanence of land-based mitigation (IPCC 2023). In this context, agroforestry has been widely incorporated into Nationally Determined Contributions as a cornerstone strategy for climate mitigation and resilience across the region. Agroforestry covers up to 357 million hectares in LAC landscapes (Somarriba et al. 2012; Krishnamurthy et al. 2019). The region also holds the world’s largest stock of tree biomass carbon on farmland (Zomer et al. 2016). Yet fewer than 15% of observations in global syntheses of soil organic carbon derive from LAC countries (Beillouin et al. 2022), and fewer than 5% of studies informing IPCC Tier-1 default coefficients for soil and biomass carbon in agroforestry were conducted in the region (Cardinael et al. 2018). Consequently, it remains unclear how well global estimates of SOC sequestration under agroforestry represent the range of conditions found across the Neotropics.

Here, we revisit the determinants of soil organic carbon sequestration in Neotropical agroforestry by quantifying the relative influence of biophysical, temporal and system descriptors. We combine three-level meta-analyses and meta-regressions on observed effect sizes, which provide the inferential basis of our conclusions, with gradient-boosted trees used to explore non-linear responses and interactions, and we assess the robustness of each finding to influential studies, apparent thresholds, confounding and data coverage. By integrating regional data, Spanish-language studies and grey literature we address four questions: (i) how much do agroforestry systems increase SOC stocks in the Neotropics; (ii) do biophysical conditions or agroforestry system characteristics explain the variability of these gains; (iii) how much of the relationship between these gains and the initial soil carbon stock follows from expressing the response relative to that stock; and (iv) how well do these drivers predict SOC responses in studies that were not used to build the models?

## 2 Materials and methods

### 2.1 Geographical scope and literature search

We systematically searched peer-reviewed and grey literature to quantify agroforestry’s impact on soil organic carbon (SOC) across a Neotropical transect (Mexico, Central America, the Caribbean, Colombia, Peru, Ecuador, and Venezuela – Fig. 2). This selection targets a high-priority bioclimatic corridor characterized by complex shaded systems and steep environmental gradients. This geographic focus leverages long-standing research partnerships (notably the AgroForesta network**-**https://agroforesta.org/), ensuring access to extensive regional grey literature. In January 2023, we searched Web of Science Core Collection, Scopus, CAB Direct, and Agricola (via OVID) supplemented by the CATIE (Centro Agronómico Tropical de Investigación y Enseñanza) institutional repository to capture regional PhD and MSc theses. Searches were conducted in English, Spanish, Portuguese, and French, provided that full texts were accessible (Table S1).

**Fig. 2.**
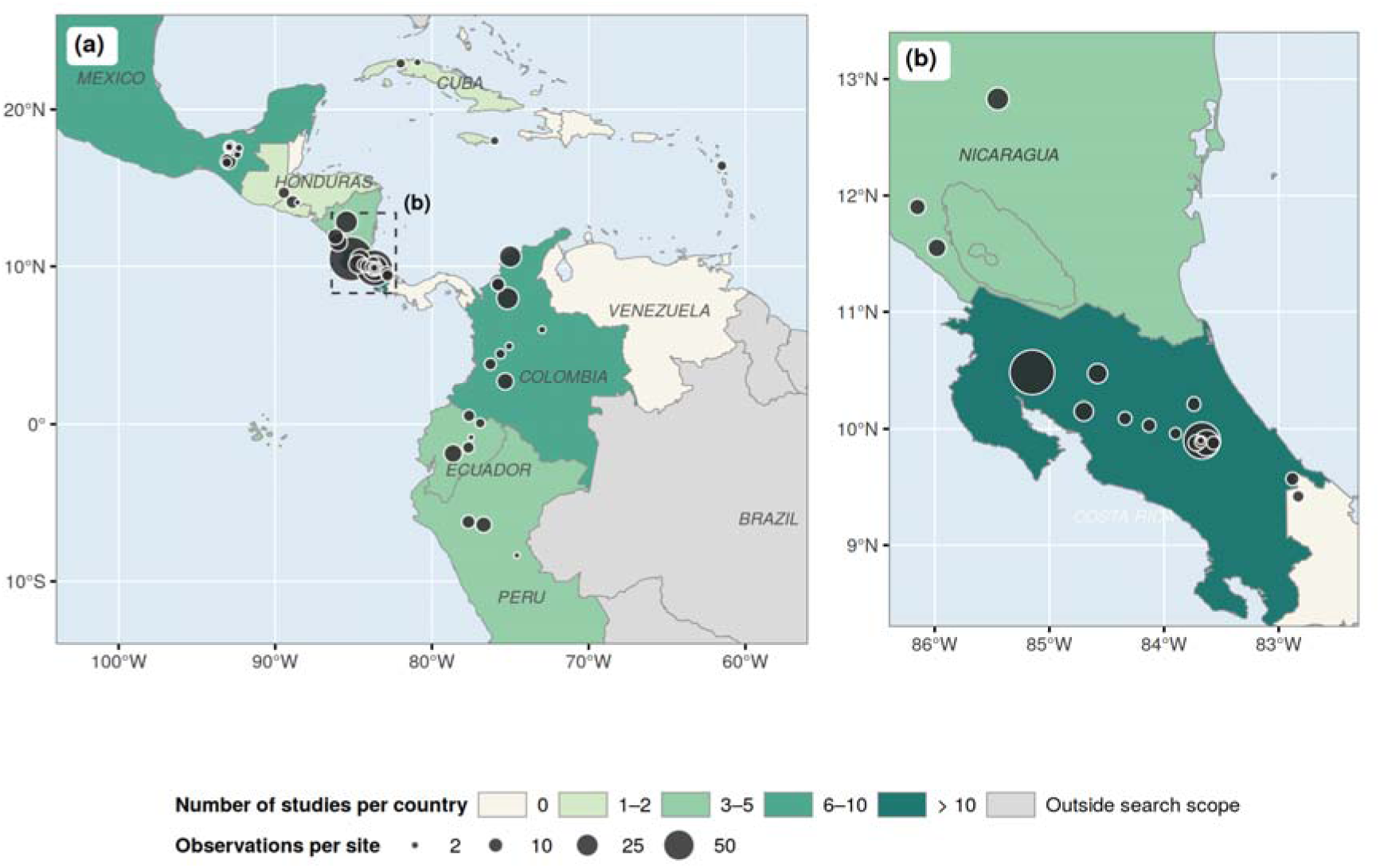
Location of the study sites. (a) Study region; the dashed rectangle delimits the area enlarged in (b). (b) Costa Rica and Nicaragua, where sites are densest. The meta-analysis includes 322 response ratios from 52 studies and 277 storage rates from 42 studies (54 studies in total). Countries within the search scope are shaded according to the number of studies, and countries outside the search scope are shown in light grey; symbol size is proportional to the number of observations at each site.

Two authors independently screened records following PRISMA guidelines, with disagreements resolved by consensus. Studies were eligible if they (i) reported field-based SOC measurements in agroforestry systems, excluding incubation or litter-only experiments; (ii) included a direct comparison with adjacent tree-less control plots (e.g., annual cropland or open-sun pasture); (iii) reported SOC as concentration (g C kg^-^¹ soil) or stock (Mg C ha^-^¹); and (iv) were conducted within the target Neotropical regions. All records were cross-checked to ensure that studies reported in both grey and peer-reviewed sources were included only once.

This screening process resulted in a final database of 322 paired observations for SOC response ratios and 277 paired observations for SOC storage rates, derived from 52 and 42 independent field studies, respectively (54 studies in total; Fig. S1). Excluded studies are listed in Table S9.

### 2.2 Data extraction

From the selected studies, we extracted mean soil organic carbon (SOC) values for agroforestry and control plots, along with sample sizes (n) and measures of dispersion (standard deviation, SD, or standard error, SE), from text, tables, or figures using WebPlotDigitizer. Grey and peer-reviewed sources were systematically cross-checked to avoid data duplication.

When SDs were not reported, they were estimated using the Extract-tool (Acutis et al. 2022) based on available SEs, confidence intervals, *t*-statistics, or *p*-values. For the remaining observations lacking any dispersion metric (24% of response ratios and 22% of storage rates), SDs were imputed using mean coefficients of variation specific to each experimental design (before–after, control–impact, or randomized designs; Fig. S2, Table S2). These coefficients were consistent with the range of SOC variability reported in independent global syntheses (Luo et al. 2006; Jian et al. 2020). Imputed standard deviations were used as estimated; a sensitivity analysis in which they were increased by 50% is reported in Table S8. When the age of the agroforestry system was not reported (56 response-ratio observations), it was set to the median age of the dataset.

For studies reporting repeated measurements over time, only the final sampling date was retained to avoid pseudoreplication and to ensure statistical independence among observations (Hurlbert 1984).

Agroforestry systems were classified into functional system types following Cardinael et al. (2018) and Nair (1991) (Table 1). Soil sampling depth was parameterized as the midpoint of the reported depth interval, a common approximation in SOC syntheses when continuous depth-resolved data are unavailable (Angers and Eriksen-Hamel 2008). Soil bulk density and coarse fragment content were extracted directly from the studies when reported; missing values were obtained from SoilGrids 2.0 at 250 m spatial resolution (Poggio et al. 2021) to allow consistent stock calculations across sites.

**Table 1.**
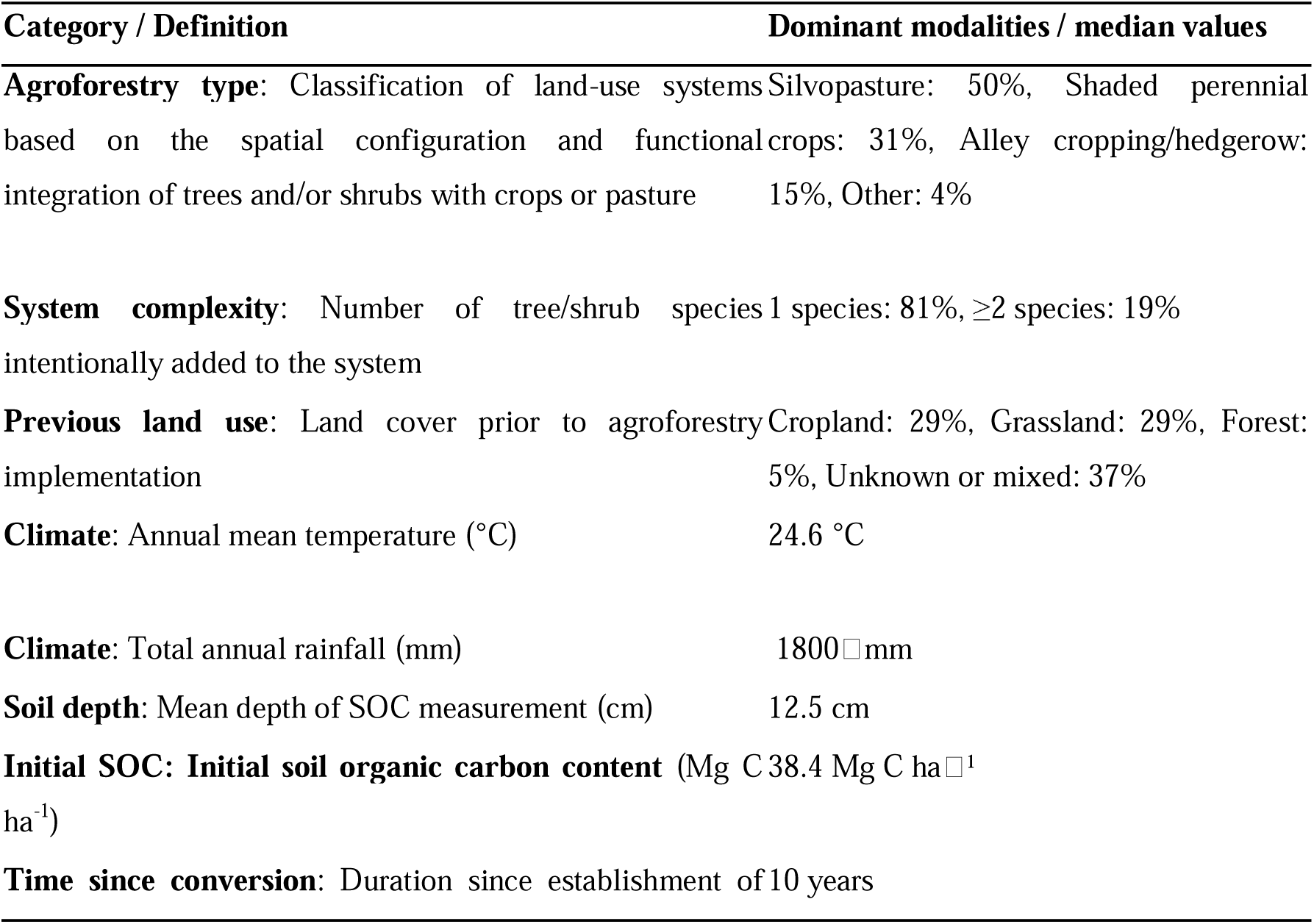
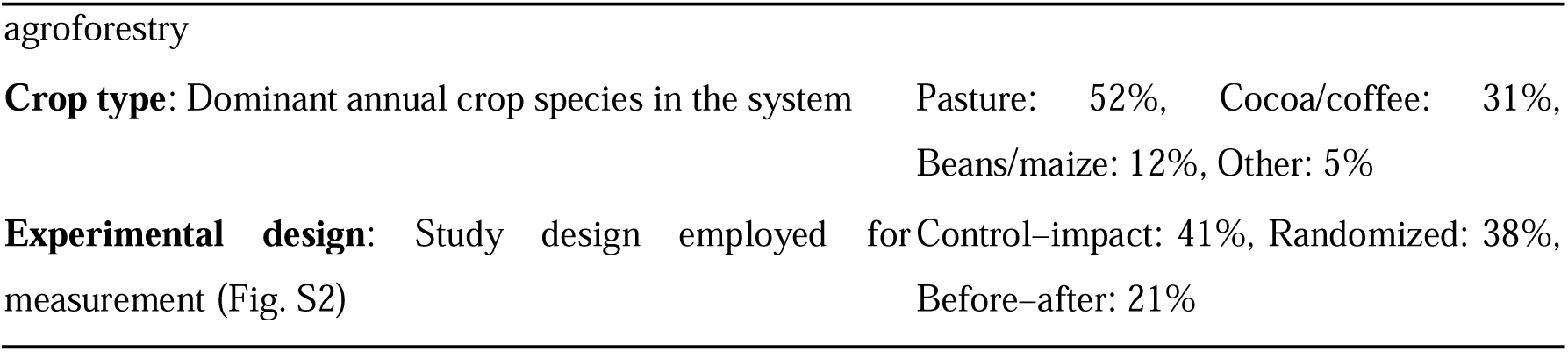
Summary of the storage-rate dataset (277 observations from 42 studies): definition of the variables and distribution of their modalities or median values. Details for both datasets are given in Table S3.

Climatic and geographic covariates, including altitude, mean annual temperature (MAT), and mean annual precipitation (MAP), were extracted from the original studies or derived from WorldClim v2.1 based on site coordinates (Fick and Hijmans 2017).

SOC stocks (Mg C ha^-^¹) were calculated or standardized when necessary. Soil organic matter values were converted to SOC using a factor of 1.72 (Nelson and Sommers 1996). SOC stocks were computed using a fixed-depth approach:

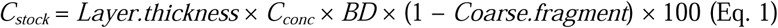

where C_stock_ is the SOC stock in Mg C ha^-^¹, C_conc_ is the SOC concentration (%), BD is the soil bulk density (g cm^-^³), Layer.thickness is soil depth interval (cm), and Coarse.fragment is the mass percentage of coarse fragments (%).

The database is available in the CIRAD Dataverse repository (https://doi.org/10.18167/DVN1/GISJUZ, version 2).

### 2.3 Standardization and effect-size calculation

The additional SOC storage (or loss) rate (*C_rate_* - Mg C ha^-^¹ yr^-^¹) was calculated for each experiment as the difference in SOC stocks changes between agroforestry and control plots, divided by the age of the agroforestry system (Eq. 2).

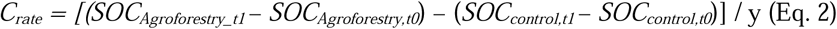

where *SOC_Agroforestry_* and *SOC_control_*represent the SOC stocks (Mg C ha^-^¹) under agroforestry and control plots, respectively, measured after (*t1*) or before (*t0*) agroforestry establishment; *y* is the agroforestry system age (years). When baseline SOC measurements were unavailable, agroforestry and control plots were assumed to have identical initial SOC stocks.

The sampling variance of C^-^rate^-^ was obtained by propagating the variances of the agroforestry and control SOC stocks, each divided by its number of replicates, and dividing the result by y² (Eq. 3):

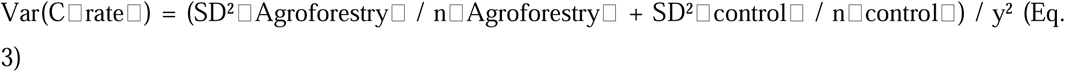

In parallel, response ratios (RR) were computed as the natural logarithm of the ratio between SOC stocks under agroforestry and control plots:

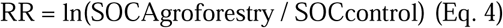

RRs and associated sampling variances were calculated using the *ROM* function in the metafor package (Viechtbauer 2010), with variances derived from large-sample approximations (Hedges et al. 1999). For ease of interpretation, log-transformed RRs were back-transformed and expressed as percentage changes in SOC (ΔCstock, %).

### 2.4 Multi-level meta-analysis and heterogeneity

To estimate overall mean effects while accounting for the non-independence of observations from the same study, we fitted three-level random-effects models, with sampling variance at the first level, variance among observations within studies at the second level and variance among studies at the third level. Likelihood ratio tests showed that both the within-study and the between-study variance components were significant for response ratios and storage rates (all p < 0.001).

The model is specified as:

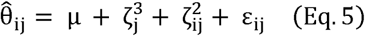

where θ^*_ij_* is the estimated effect size for the *i*-th observation in the *j*-th study, *μ*is the overall mean effect, 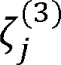 represents between-study random effects, 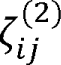 captures within-study variation among effect sizes, and ε*_ij_* is the sampling error.

Variance components were estimated by restricted maximum likelihood with the metafor package (Viechtbauer 2010). Heterogeneity was quantified with the multilevel I² statistic and partitioned between the within- and between-study levels (Nakagawa and Santos 2012). Small-study effects were evaluated with funnel plots and multilevel Egger-type regressions including the standard error of each effect size as a moderator (Egger et al. 1997), and differences between Spanish- and English-language studies and between grey literature and peer-reviewed articles were tested with likelihood ratio tests. Because the precision of SOC estimates may depend on study duration and design, residual heterogeneity was also modeled as a function of time since conversion and experimental design with location-scale models (Viechtbauer and López-López 2022).

### 2.5 Moderator analyses and robustness

The association between each moderator and the observed effect sizes was tested with three-level meta-regressions fitted by maximum likelihood and compared with the corresponding null model by likelihood ratio tests. Continuous moderators (initial SOC stock of the control, time since conversion, mean annual temperature, mean annual precipitation and sampling depth) were modeled with natural cubic splines with three degrees of freedom to allow non-linear responses, and with linear terms to estimate slopes. Categorical moderators were agroforestry type, previous land use, number of tree species added, main crop and study design. The share of heterogeneity explained by a set of moderators was computed as the proportional reduction of the summed variance components relative to the null model, both fitted by restricted maximum likelihood, for a biophysical model (initial SOC stock, time since conversion, temperature, precipitation and depth) and a management model (all categorical descriptors). Interactions between initial SOC stock and temperature, and between soil depth and each continuous moderator, were tested in the same way.

The robustness of each result was assessed in four ways. First, models were refitted after removing each study in turn; a result was considered study-dependent if the removal of a single study made it non-significant (p > 0.05) or reversed the sign of the key coefficient. Second, confounding was examined by testing each climatic effect after accounting for agroforestry type, initial SOC stock and the other climatic variable, and by comparing temperature with altitude. Third, the number of observations and of studies in each combination of depth class and time since conversion was tabulated (Fig. S7), and no prediction was made for combinations that no study sampled. Fourth, the temperature response was tested for a breakpoint by comparing linear, spline and piecewise-linear models with the Akaike information criterion, and by refitting without the sparsely sampled sites above 28 °C. Fifth, because both response variables are normalised by the control stock, the relationship between initial SOC stock and the carbon gain was repeated on the absolute difference between agroforestry and control (Mg C ha^-^¹), with time since conversion as a covariate rather than a divisor, and the resulting slope was corrected for regression to the mean following Blomqvist (1977), as implemented by Slessarev et al. (2023).

### 2.6 Machine-learning analyses

We used gradient-boosted regression trees (XGBoost; Chen et al. 2019) and a random forest (MetaForest; Van Lissa 2020) to examine non-linear responses and interactions among the same ten moderators. Their out-of-bag R² is reported in Table S5. Observations with absolute residuals greater than three standard deviations were excluded from the machine-learning analyses as influential outliers (four for response ratios and five for storage rates). XGBoost hyperparameters were fixed (maximum tree depth 4, learning rate 0.05, row subsampling 0.9, column subsampling 0.8) and the number of boosting rounds was selected by early stopping in five-fold cross-validation with folds grouped by study. Because observations from the same study are not independent, predictive performance was estimated with three cross-validation schemes, each repeated ten times: folds formed by randomly assigned observations, by soil profiles (all depths of the same agroforestry–control comparison kept together) and by studies (all observations of a study kept together). Only the last scheme measures the ability of the models to predict SOC responses in studies that were not used for training (Roberts et al. 2017).

Variable importance profiles (VIPs) were derived from gradient-boosted decision trees using the Gain metric, which quantifies the contribution of each predictor to reductions in training loss across all tree splits, summed over the indicator variables of each categorical moderator. Analyses were performed separately for response ratios and storage rates using identical predictor sets to ensure comparability. To account for sampling variability and for the nesting of observations within studies, VIPs were estimated within a bootstrap framework (Efron and Tibshirani 1993) based on 100 resamples of studies with replacement. For each resample, an XGBoost model was fitted and Gain-based importance scores were extracted. Final VIPs were obtained by averaging across bootstrap iterations, and uncertainty was quantified using the central 66% interval (17th–83rd percentiles). Interaction patterns were assessed using SHAP interaction values of the final models (Lundberg and Lee 2017), aggregated by moderator to decompose the contribution of each moderator into main and interaction components and to describe the distribution of its interactions among partner moderators.

To facilitate interpretation of model outputs, partial dependence plots (PDPs) were generated to depict marginal relationships between individual moderators and predicted SOC responses. PDPs were stratified by soil depth categories and computed, within each category, only over the range of moderator values observed in that category, to avoid extrapolation; uncertainty was estimated from 200 models refitted on bootstrap resamples of studies. These model-based curves were used solely as descriptive tools (Fig. S3); inferences on moderator effects rest on the meta-regressions described in Section 2.5.

All analyses were conducted in R version 4.4.1 with the metafor, metaforest, xgboost and ggplot2 packages. The complete workflow, from the frozen database to every number, table and figure reported in this article, is openly available (see Code availability).

## 3 Results

### 3.1 Soil carbon gains are consistent but highly heterogeneous

Across the 54 studies (Fig. 2), agroforestry increased SOC stocks by 6.8% (95% CI: 1.7–12.2%; 322 observations from 52 studies) and SOC storage rates by 0.25 Mg C ha^-^¹ yr^-^¹ (95% CI: 0.05–0.45; 277 observations from 42 studies). Estimates restricted to observations sampled within the top 30 cm were similar (6.8%, 95% CI: 1.0–13.0%; 0.24 Mg C ha^-^¹ yr^-^¹, 95% CI: 0.01 to 0.46). Heterogeneity was large (I² = 88% for response ratios and 93% for storage rates), with 67% and 75% of it, respectively, located between studies. The effect estimated from Spanish-language studies (43% of studies) did not differ from English-language ones (response ratios: likelihood ratio test (LRT) = 0.32, p = 0.57; storage rates: LRT = 0.93, p = 0.33), grey literature did not differ from peer-reviewed articles (p = 0.43 and 0.17), and multilevel Egger-type regressions detected no small-study effects (p = 0.60 and 0.54; Fig. S4).

### 3.2 Biophysical conditions, not system characteristics, explain the heterogeneity

When tested separately, initial SOC stock, time since conversion, temperature and precipitation showed significant relationships with response ratios, and initial SOC stock and temperature with storage rates. None of the system descriptors (i.e., agroforestry type, previous land use, number of tree species added, main crop and study design) was related to either response variable (all p ≥ 0.17; Table S4). Together, biophysical moderators explained 15.9% of the heterogeneity in response ratios (LRT = 43.6, df = 13, p < 0.001) and 22.7% in storage rates (LRT = 47.9, df = 13, p < 0.001), whereas system descriptors explained none (p = 0.87 and 0.77). Estimated separately within each category, mean responses ranged from +2% to +11% across agroforestry types, previous land uses and classes of added tree species, and from 0.15 to 0.44 Mg C ha^-^¹ yr^-^¹ for storage rates; all confidence intervals overlapped: no category could be distinguished from another (Table 2).

**Table 2.** Soil organic carbon responses to agroforestry by agroforestry type, previous land use and number of tree species added, estimated with three-level meta-analytic models. Response ratios are back-transformed (ratio of agroforestry to control stocks; values above 1 indicate gains) and storage rates are expressed in Mg C ha^-^¹ yr^-^¹; brackets give 95% confidence intervals. N gives the number of observations and studies. Environmental conditions are means ± standard deviations in the response-ratio dataset. Standardized XGBoost predictions are given in Table S7.

| Moderator | Level | Response ratio<br>[95% CI] | N | Storage rate (Mg C ha<sup>-1</sup><br>yr<sup>-1</sup>) [95% CI] | N | Depth (cm) | Initial SOC (Mg<br>C ha<sup>-1</sup>) | Precipitation<br>(mm) | Temperature<br>(°C) |
| --- | --- | --- | --- | --- | --- | --- | --- | --- | --- |
| Agroforestry type | Silvopasture | 1.10 [1.03; 1.17] | 172/29 | 0.23 [-0.04; 0.50] | 139/23 | 24.7 ± 17.2 | 48.2 ± 34.8 | 1941 ± 849 | 26.2 ± 3.4 |
|  | Shaded perennial crops | 1.05 [0.96; 1.15] | 72/13 | 0.29 [-0.08; 0.67] | 85/11 | 13.3 ± 13.0 | 50.8 ± 35.2 | 2538 ± 643 | 22.0 ± 2.5 |
|  | Alley<br>cropping/hedgerow | 1.04 [0.92; 1.17] | 43/8 | 0.15 [-0.26; 0.57] | 42/8 | 13.0 ± 9.3 | 58.3 ± 29.5 | 2130 ± 971 | 20.4 ± 3.5 |
|  | Multistrata | 1.02 [0.90; 1.16] | 32/8 | – | – | 22.0 ± 9.3 | 114.3 ± 74.3 | 1858 ± 673 | 25.3 ± 1.6 |
|  | Other | 1.06 [0.81; 1.39] | 3/2 | 0.44 [-0.16; 1.03] | 11/4 | 15.0 ± 0.0 | 70.9 ± 21.7 | 2353 ± 1955 | 25.6 ± 2.1 |
| Previous land use | Cropland | 1.11 [1.00; 1.22] | 75/13 | 0.18 [-0.10; 0.46] | 81/13 | 14.5 ± 12.0 | 54.8 ± 36.9 | 2078 ± 831 | 21.0 ± 3.1 |
|  | Grassland | 1.08 [0.95; 1.23] | 78/7 | 0.38 [-0.09; 0.85] | 81/8 | 23.7 ± 17.0 | 39.1 ± 22.2 | 1546 ± 426 | 26.1 ± 3.7 |
|  | Forest | 1.05 [0.89; 1.24] | 13/4 | 0.41 [-0.16; 0.97] | 13/5 | 11.5 ± 11.2 | 37.6 ± 25.6 | 2308 ± 1132 | 24.7 ± 4.0 |
|  | Unknown or mixed | 1.05 [0.98; 1.12] | 156/30 | 0.21 [-0.05; 0.47] | 102/19 | 21.9 ± 15.9 | 68.4 ± 52.6 | 2361 ± 875 | 25.1 ± 3.1 |
| Tree species added | One | 1.06 [1.01; 1.12] | 257/39 | 0.22 [0.01; 0.44] | 223/33 | 20.1 ± 15.9 | 51.6 ± 35.1 | 2210 ± 824 | 24.2 ± 3.9 |
|  | Two or more | 1.08 [1.00; 1.17] | 65/17 | 0.31 [0.04; 0.59] | 54/12 | 20.6 ± 14.5 | 77.9 ± 65.4 | 1641 ± 825 | 24.9 ± 3.4 |

Gradient-boosted trees ranked the moderators consistently for both response variables (Fig. 3a). Initial SOC stock was the most important predictor (34% of total importance for response ratios and 45% for storage rates), followed by time since conversion (15% for both), temperature (14% and 13%) and precipitation (14% and 9%), whereas each system descriptor accounted for less than 5%. Fitted to the absolute difference between agroforestry and control instead of the response ratio, the variance-weighted forest moved initial SOC stock from second to fifth in importance and time since conversion to second (Table S11). Moderators also interacted with one another, with interactions accounting for 39–75% of each moderator’s contribution to the predictions. These interactions involved several moderators rather than being concentrated on a single one (Fig. 3b, c).

**Fig. 3.**
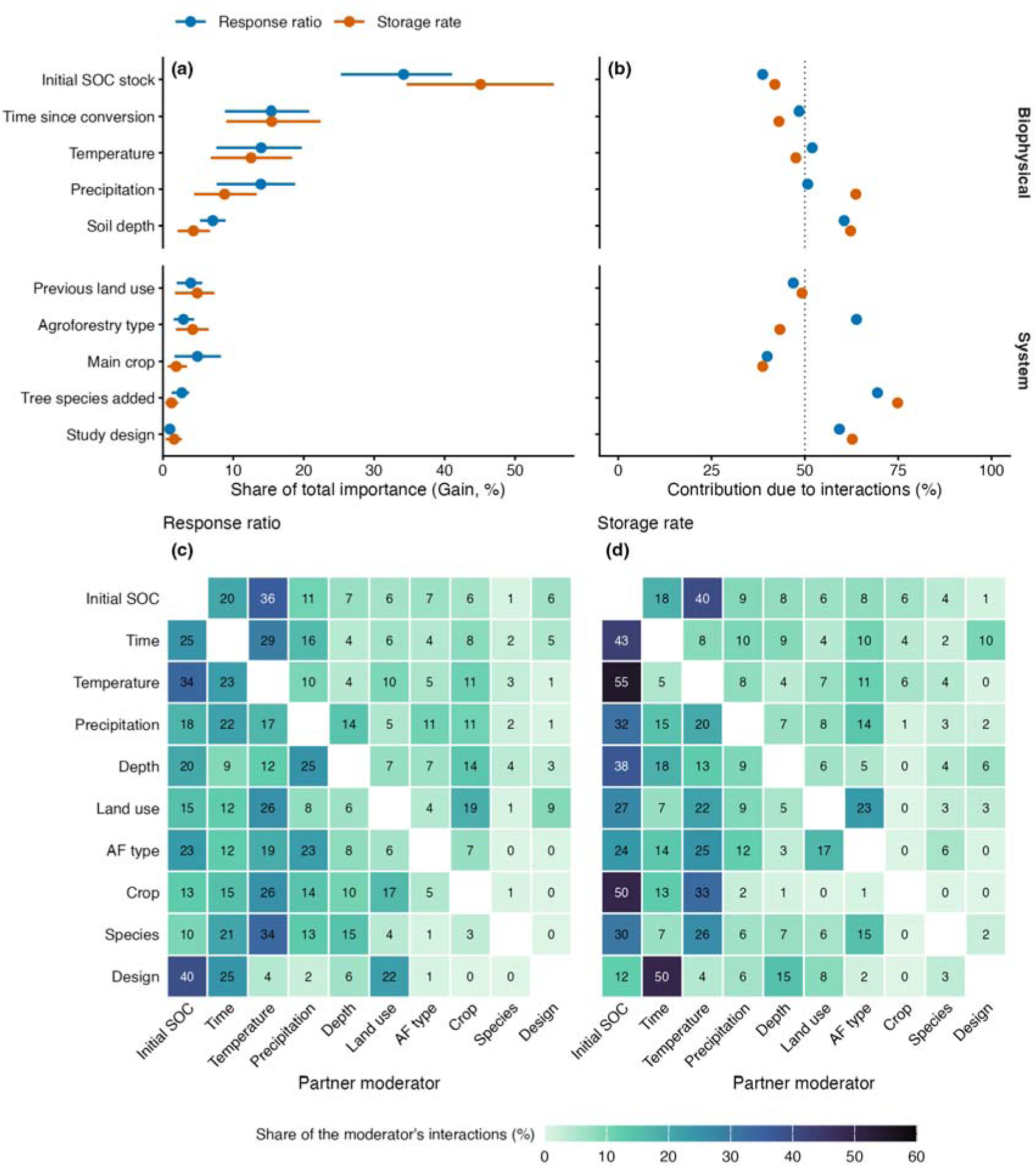
Importance and interactions of moderators in gradient-boosted tree models for response ratios and storage rates. (a) Share of total Gain importance of each moderator; points show means and lines the 17th–83rd percentiles over 100 bootstrap resamples of studies. (b) Share of the SHAP contribution of each moderator due to interactions with other moderators; the dotted line marks 50%. Moderators are grouped into biophysical conditions and system descriptors. (c, d) Distribution of the interaction contributions of each moderator (rows) among partner moderators (columns), in percent of its total interaction contribution, for (c) response ratios and (d) storage rates; values are printed in each cell. Predictive performance is reported in Table S5.

The models fitted the compiled data well (in-sample R² = 0.73 for response ratios and 0.58 for storage rates). Their performance was lower, but still appreciable, when observations were randomly divided between training and validation sets (R² = 0.49 and 0.30). It dropped further when all observations from the same soil profile were kept together during validation (R² = 0.13 and 0.08), and was close to zero when entire studies were left out of the training data (R² = 0.03 and 0.06; Table S5, Fig. S5).

### 3.3 Relative gains decline with initial soil carbon, more steeply in warm sites

Response ratios declined as initial SOC stock increased (Fig. 4a). Predicted gains were 19.2% (95% CI: 10.7–28.2%) at 20 Mg C ha^-^¹, 7.8% (1.4–14.5%) at 40 Mg C ha^-^¹, 4.5% (−1.2 to 10.5%) at 60 Mg C ha^-^¹ and 1.3% (−5.3 to 8.4%) at 90 Mg C ha^-^¹. This corresponds to a decrease of 1.8% per 10 Mg C ha^-^¹ (95% CI: 1.0–2.7; *p* < 0.001). Each initial SOC class included at least nine studies, and the relationship remained significant when studies were removed one at a time (maximum *p* = 0.001; Fig. S6, Table S6). Storage rates also decreased with initial SOC stock (−0.07 Mg C ha^-^¹ yr^-^¹ per 10 Mg C ha^-^¹; 95% CI: −0.13 to −0.01; p = 0.025). This decline was non-linear (LRT = 18.4, df = 3, p < 0.001), reaching −1.3 Mg C ha^-^¹ yr^-^¹ (−2.1 to −0.5) at 130 Mg C ha^-^¹, based on 18 observations from four studies. Unlike response ratios, the linear trend for storage rates depended on individual studies: removing any one of three studies raised its p-value above 0.05 (maximum p = 0.10). This fragility was specific to the weighting used: with the imputed standard deviations increased by 50%, the trend held after the removal of any single study (Table S8). On the absolute scale, agroforestry added 2.9 Mg C ha^-^¹ (95% CI: 0.3–5.5), and this amount did not vary with initial SOC stock (p = 0.88, with time since conversion and temperature in the model). The negative slope of the untransformed difference (−0.15 Mg C ha^-^¹ per Mg C ha^-^¹; p < 0.001) came from control–impact designs alone and disappeared after correction for regression to the mean (−0.02); it was absent in randomized designs (+0.05; p = 0.34).

**Fig. 4.**
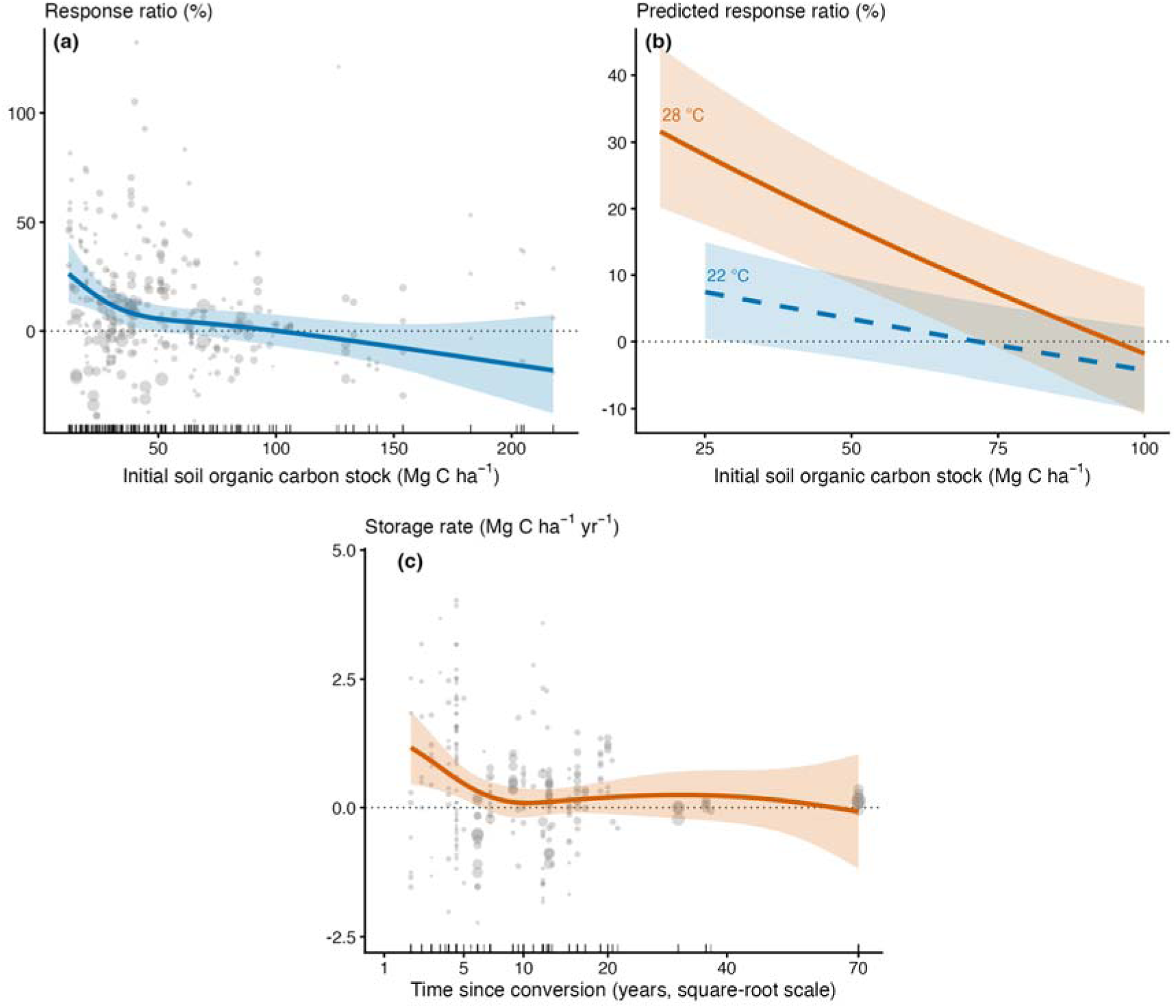
Drivers of soil organic carbon responses estimated with three-level meta-regressions on observed effect sizes. (a) Response ratio (%) as a function of the initial soil organic carbon stock of the control; the line and band show the predictions and 95% confidence intervals of a natural-spline model, points show observations sized by their precision and ticks show the distribution of observations. (b) Predicted response ratio as a function of initial soil organic carbon stock at 22 °C (dashed line) and 28 °C (solid line), each restricted to stocks between the 10th percentile observed at that temperature and 100 Mg C ha^-^¹, above which few warm sites were available. (c) Storage rate as a function of time since conversion (square-root scale). In (a) and (c), observations outside the 1st–99th percentiles are not displayed.

The effect of initial SOC stock on the potential of agroforestry to add carbon depended on temperature (LRT = 11.5, df = 1, p < 0.001; Fig. 4b). The decline in response ratios with initial SOC was steeper at 28 °C (−3.5% per 10 Mg C) than at 22 °C (−1.5% per 10 Mg C). At 30 Mg C ha^-^¹, the predicted gain was 25.8% (15.9–36.7%) at 28 °C and 6.6% (−0.1 to 13.8%) at 22 °C. At 80 Mg C ha^-^¹, neither estimate differed from zero (5.4%, −2.9 to 14.5%; −1.3%, −7.0 to 4.7%). The interaction remained significant when studies were removed one at a time (maximum p = 0.010) and after accounting for agroforestry type (*p* < 0.001). For storage rates, the slope was also more negative at 28 °C than at 22 °C (−0.09 versus −0.05 Mg C ha^-^¹ yr^-^¹ per 10 Mg C ha^-^¹), but the difference between slopes was not significant (p = 0.33). On the absolute difference between agroforestry and control, this interaction was not detectable either (LRT = 1.9, df = 1, p = 0.17; Table S10).

### 3.4 Time since conversion

Response ratios increased by 0.4% per year since conversion (95% CI: 0.04–0.8%; p = 0.037), although the relationship was non-linear (LRT = 11.7, df = 3, p = 0.009) and its shape was poorly constrained: only 24 observations from 7 studies came from systems older than 20 years. Storage rates were highest in the first years and declined thereafter: 1.17 Mg C ha^-^¹ yr^-^¹ (0.46–1.88) at 2 years, 0.47 (0.16–0.78) at 5 years, and 0.09 (−0.20 to 0.37) at 10 years, with no significant increase beyond 10 years (Fig. 4c). This decline was not statistically significant (LRT = 7.1, df = 3, p = 0.068). On the absolute difference between agroforestry and control, gains increased with time since conversion (LRT = 15.4, df = 3, p = 0.002, with initial stock and temperature in the model), and this trend held when each study was removed in turn (maximum p = 0.035; Table S10). Only 24 response ratios from 7 studies and 19 storage rates from 5 studies covered more than 20 years, and all measurements were centred within the top 25 cm of soil (Fig. S7).

### 3.5 Climate and soil depth

Temperature was associated with both response variables, although the pattern differed between them (Table S6). For response ratios, the temperature effect was no longer significant after accounting for initial SOC stock, precipitation and agroforestry type (p = 0.19). For storage rates, temperature remained significant after accounting for these variables (p < 0.001), after removing each study in turn (maximum p = 0.006), and after excluding the warmest sites (p = 0.014). The relationship was not linear: predicted storage rates showed no consistent trend between 18 and 27 °C and were highest at 29 °C (1.33 Mg C ha^-^¹ yr^-^¹; 95% CI: 0.74–1.92), but this estimate was based on four observations from a single study. A piecewise model with a break at 27 °C had a lower AIC than the linear model (ΔAIC = 5.4), but the break was just below the temperature of that study. The apparent change around 27 °C should not be interpreted as a thermal threshold. Temperature was also strongly correlated with altitude in both datasets (r = −0.87 and −0.89), making it difficult to separate their effects: altitude fitted response ratios slightly better than temperature (AIC lower by 2.3), whereas temperature fitted storage rates clearly better (AIC lower by 7.5). These results globally suggest that temperature is related to SOC storage rates, but the form of this relationship remains uncertain.

Soil depth was not associated with response ratios (p = 0.60) or storage rates (p = 0.60), and did not change the effects of initial SOC stock, time since conversion or temperature (all *p* > 0.4). Depth was entered as the midpoint of the sampled interval, meaning that the same midpoint could represent layers of very different thicknesses (median 20 cm, range 3.5–100 cm). Adding layer thickness to the models did not change the results: depth remained non-significant (p = 0.35 and 0.42), and thickness was not related to either response variable (p = 0.55 and 0.85). Restricting the analysis to surface layers also gave the same result (216 observations from 52 studies for response ratios and 166 from 41 studies for storage rates; all p ≥ 0.21).

The depth × precipitation interaction was significant for response ratios (LRT = 10.9, df = 1, *p* < 0.001) but not for storage rates (p = 0.19). For response ratios, the interaction remained significant when studies were removed one at a time (maximum *p* = 0.034). At 50 cm, predicted gains were 28.3% (13.0–45.7%) at 1200 mm and −2.1% (−10.9 to 7.7%) at 2800 mm. At 10 cm, the corresponding estimates were 5.9% (−0.9 to 13.2%) and 6.9% (0.9–13.2%). The interaction was based on 99 observations sampled at 30 cm or deeper from 16 studies, with system ages ranging from 3 to 19 years. For response ratios, precipitation was not significant after accounting for the other biophysical moderators (p = 0.20).

### 3.6 Precision of estimates depends on study design and duration

Residual heterogeneity depended on experimental design and on time since conversion (location-scale models: p < 0.001 for both response variables; Fig. S8). For storage rates, the standard deviation of true effects fell by 31% between 2 and 10 years after conversion; for response ratios it rose by 14%. The two metrics differ by construction here: a rate is a stock change divided by a duration, so a longer duration compresses both the estimate and its spread, whereas a ratio involves no such division. At a given age, randomized designs were far more variable than before–after designs, with standard deviations larger by 98% for storage rates and 61% for response ratios. Design therefore changed how precisely an effect could be estimated, not how large it was: mean responses did not differ between designs (p = 0.63 for response ratios and 0.18 for storage rates).

## 4 Discussion

### 4.1 Initial soil carbon stock sets the relative gain, not the amount stored

Initial SOC stock was the strongest predictor in both sets of models: relative gains and storage rates declined as initial SOC increased, with storage rates becoming negative at the highest stocks. The decline in relative gains was robust to the removal of individual studies, whereas the decline in storage rates was not. This pattern can arise because the control stock is part of both the response ratio and its explanatory variable. Random error in the control measurement can then induce a negative relationship: underestimating the control stock inflates the apparent gain, whereas overestimating it reduces the gain (Slessarev et al. 2023). We therefore tested the absolute difference between agroforestry and control, which does not scale the response by the control stock. The relationship disappeared, in the meta-regressions as in the machine-learning ranking (Table S11). The amount of carbon added did not vary with initial stock, and the negative slope in the untransformed difference was driven by control–impact designs and disappeared after correcting for regression to the mean (Section 3.3, Table S10). The decline in relative gains therefore reflects the denominator: the same addition represents a smaller fraction of a larger stock. Its temperature dependence was also not detectable on the absolute scale (p = 0.17). We therefore interpret the steeper decline at 28 °C as a feature of relative gains, not as evidence for a temperature effect on carbon storage. Storage rates are subject to the same dependence on the control stock, since it enters their numerator, and are additionally divided by the duration of the comparison. Their decline was consequently the least stable of the relationships we examined.

The decline with initial stock is often interpreted as evidence of soil carbon saturation, whereby the capacity of mineral surfaces to stabilise organic matter becomes increasingly limited as carbon accumulates (Six et al. 2002; Stewart et al. 2007; Georgiou et al. 2022). We find no evidence for this in our data: absolute carbon gains did not decline with initial stock. A direct test of saturation would require information on the fine fraction and on the distribution of carbon among soil pools (Hassink 1997; Cotrufo et al. 2019), which was rarely available in the primary studies.

Carbon added by the same intervention can therefore represent a larger relative gain in a carbon-poor soil without implying that more carbon is stored there. Targeting carbon-depleted soils may thus produce larger improvements in soil condition without increasing the amount of carbon added (Post and Kwon 2000; Sanderman et al. 2017; Minasny et al. 2017). The distinction also matters when baseline stocks are used to predict carbon gains, as in crediting schemes or models calibrated on published syntheses. Because relative changes are mathematically dependent on the initial stock, part of the apparent relationship can arise from this scaling rather than from differences in how soils respond to the intervention. Reporting absolute changes alongside relative changes would allow these effects to be distinguished in primary studies.

### 4.2 Gains accumulate during the first years after establishment

Relative gains increased with time since conversion. The absolute amount of carbon added increased as well, but not in proportion to time (LRT = 15.4, df = 3, p = 0.002; Section 3.4, Table S10), and this result was unchanged when individual studies were removed. Storage rates showed the opposite pattern: they were highest soon after conversion and were no longer distinguishable from zero after 10 years, although the decline was not significant (p = 0.068). A similar decline in sequestration rates with stand age was reported by Kim et al. (2016) in a global compilation. These patterns are compatible because the storage rate is the accumulated difference in stocks divided by the time elapsed. It can therefore decline while the difference in stocks continues to increase. A declining rate does not by itself indicate that accumulation has stopped. A single annual rate is consequently a poor descriptor of the carbon accumulation trajectory. Its value depends on the age of the system when it was measured and on the period over which the change was calculated, so the timescale to which a rate applies needs to be stated explicitly (Don et al. 2024). Some of the variation among published sequestration rates may therefore reflect when systems were measured rather than differences among systems themselves.

These patterns have large implications for projections of agroforestry sequestration potential. Large-scale assessments have taken different approaches. Some apply a fixed annual rate to an area. For example, Bossio et al. (2020) use rates of 0.59 and 0.69 Mg C ha^-^¹ yr^-^¹ for alley cropping and windbreaks, respectively, based on a literature review, while noting that accumulation should eventually slow but without representing this explicitly in the calculation. Others annualise a stock difference over the 20-year transition period used in IPCC guidance (IPCC 1997, 2019). Roe et al. (2021), for example, estimate soil carbon change on cropland and grassland as the difference between projected 2038 and 2018 stocks divided by 20 years, while soil-carbon crediting methodologies similarly spread a stock change linearly over the crediting period (Gold Standard 2020). A third approach is to model accumulation as a saturating trajectory. Sommer and Bossio (2014) described sequestration using a sigmoid function, and Zomer et al. (2017) applied a similar approach to global cropland while restricting the projection to its approximately linear first 20 years. Because accumulation is expected to slow as soils approach a new steady state (West and Six 2007; Don et al. 2024), this latter approach is more consistent with the temporal pattern observed here. To our knowledge, however, no large-scale assessment has estimated an accumulation trajectory explicitly as a function of time since conversion in agroforestry systems. This is the relationship that would be needed to project carbon accumulation from the temporal dependence observed here. This has proven difficult, because long-term evidence remains limited. Only 24 observations from 7 studies for relative gains and 19 from 5 studies for storage rates extended beyond 20 years, and all were centred within the top 25 cm.

### 4.3 Climate modulates the response, and its effect depends on sampling depth

The temperature effect on relative gains was no longer significant after accounting for other covariates. It persisted for storage rates, although the fitted relationship was not monotonic and its warm end was based on only four observations from a single site. Faster organic matter turnover in warmer soils may be offset by greater plant carbon inputs, weakening the direct temperature signal (Davidson and Janssens 2006; Anderson-Teixeira et al. 2016). Larger effects have been reported in arid and seasonally dry environments (Pan et al. 2025; Ngaba et al. 2024), although these syntheses classify climate rather than testing continuous gradients.

Precipitation affected the response through sampling depth: the agroforestry–control contrast increased with depth at the drier end of the gradient, but not at the wetter end. The pattern did not rest on a single study, and deeper rooting under water limitation provides a plausible explanation (Schenk and Jackson 2002). If this pattern is confirmed, a fixed 0–30 cm sampling depth could give climate-dependent estimates of the agroforestry effect by missing a larger share of the response in drier systems. How far it extends into genuinely dry climates cannot be established here, with only 17 observations from 4 studies below 1000 mm and no deep observations from systems older than 19 years. More depth-resolved measurements in drier systems are needed.

Apart from this interaction, sampling depth was unrelated to either response variable. This result was unchanged when layer thickness was accounted for and when the analysis was restricted to layers starting at the surface. It differs from studies reporting lower sequestration rates at depth, such as Hübner et al. (2021), who found rates declining from 0.92 to 0.52 Mg C ha^-^¹ yr^-^¹ between 0–20 and 40–60 cm in Chinese agroforestry. The difference likely reflects what is being measured: absolute carbon accumulation generally decreases with depth because deeper layers contain less carbon and receive smaller inputs, whereas our response variables quantify the difference between agroforestry and control at the same depth. The absence of a depth gradient in the relative response indicates that the agroforestry–control contrast does not necessarily weaken with depth. Whether this reflects deeper root inputs, downward movement of dissolved organic carbon, or changes in belowground turnover cannot be resolved from the available data. This limits our ability to assess the mechanisms of subsoil carbon change and whether deeper layers contribute additional sequestration (Poeplau and Don 2013), particularly given the concentration of observations in relatively young systems and shallow layers.

### 4.4 System type carries no information on the soil carbon response

Agroforestry and soil-carbon syntheses commonly organise their results by system type (De Stefano and Jacobson 2018; Hübner et al. 2021; Pan et al. 2025), as do carbon-accounting frameworks when assigning default values (IPCC 2019; Cardinael et al. 2018). In our dataset, agroforestry type, tree species added, previous land use, main crop and study design explained none of the heterogeneity in either response variable, whereas the biophysical moderators did. Estimates for the individual categories overlapped widely (Table 2), so the data exclude large systematic differences between categories without excluding small ones. This differs from the result of Ma et al. (2020), who reported an effect of tree species diversity on carbon accumulation in agroforestry systems. Their estimate concerns total ecosystem carbon, including biomass, whereas ours concerns soil carbon, and our species descriptor separates broad classes of trees added and does not measure diversity itself. The two results do not necessarily refer to the same process. Tree diversity may affect carbon accumulation through biomass production without producing a detectable difference in soil carbon, or a broader diversity gradient may be needed to detect such an effect. Our descriptor also counts species without identifying them, so it does not separate nitrogen-fixing trees, which dominate the shade canopy of Neotropical coffee and cacao systems. A null result on the number of species therefore does not exclude an effect of species identity.

The categories themselves also combine systems that differ in tree density, species composition and structure (Hübner et al. 2021; Terasaki Hart et al. 2023). They are not independent of climate either: mean annual temperature ranged from 20.4 °C in alley cropping and hedgerows to 26.2 °C in silvopastures, which account for more than half of our observations and for most of those above 26 °C. A difference between system types can therefore partly reflect where those systems are established. The same problem occurs in the synthesis literature, where results are often reported for unspecified agroforestry systems rather than separated by system type (Schievano et al. 2026), making type-specific comparisons difficult.

The absence of a system-type effect does not mean that management is unimportant. Tree density, pruning and residue management were too rarely reported in the primary studies to be included in the analysis. These variables may explain substantial variation within the broad categories used here, but this could not be tested with the available data.

### 4.5 Comparison with published values and implications for carbon accounting

The gains estimated here are lower than most previously published values. Relative to tree-less agricultural controls, previous syntheses reported SOC increases of 19% over 0–100 cm (Shi et al. 2018), 26–40% depending on soil depth (De Stefano and Jacobson 2018), and 20% (95% CI, 17–23%) in a synthesis of 230 meta-analyses (Beillouin et al. 2023), although the most recent global synthesis reports 10.7% against controls that include forest (Pan et al. 2025). The IPCC Tier 1 stock-change factor for cropland converted to agroforestry is 1.25 ± 0.04 over 0–30 cm (Cardinael et al. 2018). Published accumulation rates are also higher, from 0.75 ± 0.19 Mg C ha^-^¹ yr^-^¹ globally for cropland converted to agroforestry (Cardinael et al. 2018) to 2.23 Mg C ha^-^¹ yr^-^¹ in tropical climates (Feliciano et al. 2018).

The difference partly reflects what each synthesis represents. Our estimate pools systems of all ages, whereas the Tier 1 factor represents a 20-year transition. At 20 years, the relative gain predicted by our model is 14.2% (5.1–24.1), closer to published values, although this estimate is based on 30 observations from 9 studies at or beyond that age. The control also matters: silvopastoral systems compared with open pasture account for 53% of our response ratios, and syntheses distinguishing control types report smaller gains over pasture than over cropland (+9 to +10% versus +26 to +40%; De Stefano and Jacobson 2018). The rate reported by Hübner et al. (2021) for the uppermost layer of silvopastoral systems, 0.23 Mg C ha^-^¹ yr^-^¹, is close to our overall estimate. Our three-level models also give less weight to studies reporting many soil layers than analyses treating observations as independent. Language and document type did not explain the discrepancy.

The implications for carbon accounting are more fundamental. IPCC Tier 1 and many Tier 2 approaches assign fixed values according to system type or land-use class, whereas system type explained none of the heterogeneity in our data and initial SOC stock and climate did. The Tier 1 factor is also multiplicative: applying a relative change to a reference stock makes the credited amount increase with that stock, whereas the absolute amount added did not increase with initial stock in our data. Accounting based on absolute change, parameterised by site conditions and time since conversion rather than system type (Section 4.2), would be more consistent with the patterns observed here (Paustian et al. 2016; Amelung et al. 2020; Sanderman et al. 2017).

### 4.6 Apparent predictive performance depends on the validation scheme

The three-level meta-regressions provide the basis for inference, while gradient-boosted trees were used to examine non-linear responses and interactions. Both approaches ranked the moderators similarly, but the predictive performance of the trees changed sharply with the validation scheme. Cross-validated R² was about 0.5 for relative gains and 0.3 for storage rates when observations from the same study could occur in both training and test folds, but fell to 0.03 and 0.06 when entire studies were withheld. Profile-level validation was intermediate (0.13 and 0.08), but its uncertainty overlapped with both levels. The contrast indicates that much of the apparent predictive power comes from information shared within studies rather than from relationships that transfer across studies. This matters for soil-carbon syntheses, where several observations often describe the same profile, site or study (Roberts et al. 2017). Study-level validation is consequently the more relevant test of prediction for a new site.

Machine-learning approaches are increasingly used in soil-carbon research (Wadoux et al. 2020; Song et al. 2025), but validation that allows related observations to occur in both training and test sets can substantially overstate performance on new sites (Stockmann et al. 2013; Reichstein et al. 2019). In our data, the trees were useful for describing non-linearities and interactions among the observed studies, but had little ability to predict responses in studies not represented in the training data. This distinction is important when meta-analytic machine learning is presented as predictive modelling: strong cross-validation within a study set does not necessarily imply transfer to new sites.

Study design does not appear to be a major source of systematic bias in the estimated agroforestry effect, but it affects the precision of the estimates: at a given age, randomized comparisons were far more variable than before–after designs. Precision also depends on duration, the standard deviation of true storage rates falling by 31% between 2 and 10 years. Short-term rates are therefore estimated from the noisiest part of the evidence, which adds to the uncertainty around using them to infer longer-term carbon accumulation.

### 4.7 Limitations and perspectives

The main limitations concern the coverage of the evidence base and the way soil carbon was measured and reported. The scarcity of observations beyond 20 years and below 30 cm limits our ability to assess whether the temporal patterns observed here persist over longer periods or extend into subsoils. The strong correlation between temperature and altitude also prevents these gradients from being separated, leaving their individual contributions uncertain.

Comparisons of soil stocks are subject to additional measurement constraints. Fixed-depth calculations can bias agroforestry–control differences when bulk density differs between systems, particularly in silvopastoral comparisons where grazing can compact the control soil. Because bulk density was incompletely reported, we could not correct stocks on an equivalent-soil-mass basis. Sampling position relative to trees was also rarely reported. This is potentially important because soil carbon can vary strongly with distance from tree rows or trunks, and sampling under trees, between rows or across the whole plot may differ systematically among systems. Such differences could contribute to the heterogeneity that our moderators did not explain. Missing dispersion measures and uncertainty in short-term storage rates add further uncertainty, particularly for studies where changes in stock are small.

These limitations point to specific priorities for future syntheses and field studies. Continuous environmental variables should be retained rather than reduced to broad system categories, but this needs to be accompanied by better coverage of poorly sampled parts of the environmental space. Long-term measurements with depth-resolved sampling are needed to establish whether carbon accumulation continues beyond the periods represented here and whether subsoils contribute substantially to the total response. Primary studies should also report sampling position, bulk density, tree density, pruning and residue management consistently. This would allow variation currently attributed to unexplained heterogeneity or system type to be separated into environmental and management effects. More complete reporting would also make it easier to combine regional evidence, including grey literature and Spanish-language studies, with the datasets used in global syntheses.

## 5 Conclusion

Agroforestry increased soil organic carbon by 0.25 Mg C ha^-^¹ yr^-^¹, or about 7%, across 54 Neotropical studies. The increase varied widely, but not according to the categories commonly used to describe agroforestry: agroforestry type, previous land use, number of tree species, main crop and study design explained none of the heterogeneity, whereas initial soil carbon, time since conversion and climate did. Here we show that the relative size of the gain is set by the state of the soil before conversion and by the climate of the site, whereas the amount of carbon added depends mainly on how long the system has been in place.

For carbon accounting, gains would be better estimated from site-specific baseline stocks, climate and system age than from generic rates or system classes. This also means that where agroforestry is established matters: targeting carbon-depleted soils maximises relative improvement, although not the quantity of carbon stored. The temporal dynamics further caution against using short-term rates for long-term accounting, as applying a rate measured over two years across a twenty-year crediting period would overestimate cumulative gains several-fold. More broadly, the limited ability of models to predict at unseen sites shows that good fit to compiled data does not necessarily translate into reliable large-scale predictions, underscoring the need for independent validation.

## Supporting information

supp Mat

## Acknowledgments

We greatly acknowledge the researchers whose published data were used in this meta-analysis; their studies are listed in the References of the meta-analysis.

## Declarations

### Ethics approval

Not applicable.

### Consent to participate

Not applicable.

### Consent for publication

Not applicable.

### Use of artificial intelligence tools

During the revision, the authors used Claude (Anthropic) to assist with code review, re-analysis scripting, figure preparation and language editing. The authors checked all analyses, outputs and text and take full responsibility for the content of this article.

### Availability of data and material

The corrected database and the two analysis datasets analyzed in this study are available in the CIRAD Dataverse repository (https://doi.org/10.18167/DVN1/GISJUZ, version 2; Demenois et al. 2025); the uncorrected database remains available as version 1. The documented data corrections, the excluded studies and the observations excluded from the machine-learning analyses are provided with the code (see Code availability); the excluded studies are listed in Table S9.

### Code availability

The R code reproducing all analyses, tables and figures from the frozen database is available at https://github.com/dbeillouin/MACCA (release v1.4-R3; Beillouin 2026) and is archived in Software Heritage (swh:1:rev:27c78e8a818fa0ab487f458f182427e3a26489c5).

### Authors’ contributions

Conceptualization, DB, RC, JD; Methodology, DB, RC, JD; Software, DB; Validation, DB, RC, JD; Formal analysis, DB; Investigation, DB, CV; Data curation, DB, CV, RC, UC, JBL, JD; Writing – Original Draft, DB, CV, RC, UC, JD; Writing – Review & Editing, DB, CV, RC, UC, JBL, PAW, JD; Visualization, DB; Supervision, JD; Project administration, JD; Funding acquisition, JD.

