## Supplementary material for "Baseline soil carbon, time since conversion and climate, not system characteristics, shape soil carbon responses in Neotropical agroforestry. A meta-analysis": supp Mat

**Electronic supplementary material**

*Agronomy for Sustainable Development*

This file contains Tables S1–S7 and Figs. S1–S8. All values were produced by the R workflow available at https://github.com/dbeillouin/MACCA from the database deposited at https://doi.org/10.18167/DVN1/GISJUZ.

**Table S1** Search string applied to titles, abstracts and keywords in Web of Science Core Collection, Scopus, CAB Direct and Agricola (via OVID) in January 2023. Searches were complemented by the CATIE institutional repository to capture theses and reports.

Search string

Soil carbon OR Soil organic matter OR SOM OR Soil organic carbon OR SOC OR Soil AND (Carbon pool* OR Carbon stock* OR Organic carbon OR Carbon OR Carbon sequestration OR Carbon concentration OR Carbon content OR Organic matter) AND (Agroforest* OR Alley cropping OR Hedgerow* OR Multistrata system* OR Agroforest* fallow* OR Parkland* OR Shaded perennial-crop system* OR Silvo-arable system* OR Silvo*pasture* OR Homegarden* OR Windbreak* OR Shelterbelt* OR live fence* OR Tree intercrop*) AND (Central America OR Panama OR Costa Rica OR Honduras OR Nicaragua OR El Salvador OR Guatemala OR Belize OR Mexico OR Yucatan OR Chiapas OR Quintana Roo OR Tabasco OR Campeche OR Caribbean OR Antilles OR Cayman Islands OR Cuba OR Jamaica OR Bahamas OR Turks and Caicos Islands OR Haiti OR Dominican Republic OR Puerto Rico OR Virgin Islands OR Anguilla OR St Martin OR Antigua and Barbuda OR Aruba OR Barbados OR Bonaire OR Curaçao OR Dominica OR Federal Dependencies of Venezuela OR Grenada OR Guadeloupe OR Martinique OR Montserrat OR Nueva Esparta OR Saba OR Archipelago of San Andres OR Saint Barthélémy OR Saint Kitts and Nevis OR Saint Lucia OR Saint Martin OR Saint Vincent and the Grenadines OR Sint Eustatius OR Sint Maarten OR Trinidad and Tobago OR French West Indies OR FWI OR Venezuela OR Bolivarian Republic of Venezuela OR Colombia OR Republic of Colombia OR Ecuador OR Republic of Ecuador OR Peru OR Republic of Peru)

**Table S2** Mean coefficients of variation (CV) of soil organic carbon stocks used to impute missing standard deviations, by experimental design. For storage rates, CVs include the 50% increase described in Section 2.2. N: observations; Imputed: observations with at least one imputed standard deviation.

| Dataset | Design | N | Imputed | CV agroforestry | CV control |
| --- | --- | --- | --- | --- | --- |
| Response ratios | Before–after | 39 | 24 | 0.116 | 0.107 |
| Response ratios | Control–impact | 177 | 32 | 0.289 | 0.286 |
| Response ratios | Randomized | 106 | 22 | 0.170 | 0.226 |
| Storage rates | Before–after | 57 | 23 | 0.181 | 0.239 |
| Storage rates | Control–impact | 114 | 15 | 0.372 | 0.370 |
| Storage rates | Randomized | 106 | 22 | 0.256 | 0.339 |

**Table S3** Characteristics of the response-ratio and storage-rate datasets. Continuous variables: median (interquartile range); range.

| Variable | Response ratios | Storage rates |
| --- | --- | --- |
| Observations / studies | 322 / 52 | 277 / 42 |
| Agroforestry type | Silvopasture 53%, shaded perennial crops 22%, alley cropping/hedgerow 13%, multistrata 10%, other 1% | Silvopasture 50%, shaded perennial crops 31%, alley cropping/hedgerow 15%, other 4% |
| Tree species added | One 80%, two or more 20% | One 81%, two or more 19% |
| Previous land use | Grassland 24%, cropland 23%, forest 4%, unknown or mixed 48% | Cropland 29%, grassland 29%, forest 5%, unknown or mixed 37% |
| Main crop | Pasture 55%, cocoa/coffee 26%, beans/maize 10%, other 9% | Pasture 52%, cocoa/coffee 31%, beans/maize 12%, other 5% |
| Study design | Control–impact 55%, randomized 33%, before–after 12% | Control–impact 41%, randomized 38%, before–after 21% |
| Mean annual temperature (°C) | 25.0 (22.0–27.2); 13–30 | 24.6 (21.8–27.2); 14–30 |
| Mean annual precipitation (mm) | 1985 (1500–2648); 95–4380 | 1800 (1500–2640); 95–4380 |
| Sampling depth, midpoint (cm) | 15 (10–30); 1.8–70 | 12.5 (7.5–30); 1.8–70 |
| Initial SOC stock (Mg C ha⁻¹) | 41.9 (27.7–71.3); 12–218 | 38.4 (26.0–63.1); 12–154 |
| Time since conversion (years) | 9 (4.5–12.7); 2–70 | 10 (4.5–12.7); 2–70 |

Table S4 Tests of moderators on observed effect sizes (three-level meta-regressions fitted by maximum likelihood; likelihood ratio tests against the null model). Heterogeneity explained is the proportional reduction of the summed variance components, truncated at zero.

| **Response** | **Moderator** | **Observations / studies** | **df** | **LRT** | **p** | **Slope [95% CI]** | **Heterogeneity explained (%)** |
| --- | --- | --- | --- | --- | --- | --- | --- |
| Response ratio | Initial SOC stock (linear) | 322 / 52 | 1 | 17.62 | < 0.001 | −0.0019 [−0.0027; −0.0010] | 0.0 |
| Response ratio | Initial SOC stock (spline) | 322 / 52 | 3 | 20.79 | < 0.001 | – | 0.0 |
| Response ratio | Time since conversion (linear) | 322 / 52 | 1 | 4.35 | 0.037 | 0.0041 [0.0004; 0.0077] | 7.3 |
| Response ratio | Time since conversion (spline) | 322 / 52 | 3 | 11.68 | 0.009 | – | 8.9 |
| Response ratio | Mean annual precipitation (spline) | 322 / 52 | 3 | 9.55 | 0.023 | – | 4.8 |
| Response ratio | Mean annual temperature (spline) | 322 / 52 | 3 | 11.56 | 0.009 | – | 6.9 |
| Response ratio | Sampling depth (spline) | 322 / 52 | 3 | 1.85 | 0.60 | – | 0.0 |
| Response ratio | Sampling depth (linear) | 322 / 52 | 1 | 0.16 | 0.69 | 0.0003 [−0.0013; 0.0020] | 0.0 |
| Response ratio | Agroforestry type | 322 / 52 | 4 | 1.68 | 0.79 | – | 0.0 |
| Response ratio | Previous land use | 322 / 52 | 3 | 1.01 | 0.80 | – | 0.0 |
| Response ratio | Tree species added | 322 / 52 | 1 | 0.21 | 0.64 | – | 0.0 |
| Response ratio | Main crop | 322 / 52 | 3 | 4.80 | 0.19 | – | 0.0 |
| Response ratio | Study design | 322 / 52 | 2 | 0.91 | 0.63 | – | 0.0 |
| Response ratio | All biophysical moderators | 322 / 52 | 13 | 43.57 | < 0.001 | – | 15.9 |
| Response ratio | All system descriptors | 322 / 52 | 13 | 7.59 | 0.87 | – | 0.0 |
| Storage rate | Initial SOC stock (linear) | 253 / 41 | 1 | 5.00 | 0.025 | −0.0068 [−0.0128; −0.0009] | 3.9 |
| Storage rate | Initial SOC stock (spline) | 253 / 41 | 3 | 18.38 | < 0.001 | – | 0.0 |
| Storage rate | Time since conversion (linear) | 253 / 41 | 1 | 1.08 | 0.30 | −0.0079 [−0.0228; 0.0070] | 0.0 |
| Storage rate | Time since conversion (spline) | 253 / 41 | 3 | 7.12 | 0.068 | – | 0.0 |
| Storage rate | Mean annual precipitation (spline) | 253 / 41 | 3 | 5.60 | 0.13 | – | 0.0 |
| Storage rate | Mean annual temperature (spline) | 253 / 41 | 3 | 17.39 | < 0.001 | – | 19.5 |
| Storage rate | Sampling depth (spline) | 253 / 41 | 3 | 1.88 | 0.60 | – | 2.3 |
| Storage rate | Sampling depth (linear) | 253 / 41 | 1 | 0.66 | 0.42 | 0.0034 [−0.0047; 0.0114] | 1.7 |
| Storage rate | Agroforestry type | 253 / 41 | 3 | 0.65 | 0.89 | – | 0.0 |
| Storage rate | Previous land use | 253 / 41 | 3 | 1.34 | 0.72 | – | 0.0 |
| Storage rate | Tree species added | 253 / 41 | 1 | 0.03 | 0.87 | – | 0.0 |
| Storage rate | Main crop | 253 / 41 | 3 | 1.36 | 0.71 | – | 0.0 |
| Storage rate | Study design | 253 / 41 | 2 | 3.46 | 0.18 | – | 3.2 |
| Storage rate | All biophysical moderators | 253 / 41 | 13 | 47.88 | < 0.001 | – | 22.7 |
| Storage rate | All system descriptors | 253 / 41 | 12 | 8.17 | 0.77 | – | 0.0 |

Table S5 Performance of the gradient-boosted trees (mean ± standard deviation over ten repetitions of five-fold cross-validation) and out-of-bag performance of the meta-analytic random forest.

| **Metric** | **Response ratios** | **Storage rates** |
| --- | --- | --- |
| Observations used (after outliers) | 318 | 248 |
| Boosting rounds (early stopping) | 76 | 32 |
| In-sample R² | 0.73 | 0.58 |
| Cross-validated R², random observations | 0.49 ± 0.02 | 0.30 ± 0.04 |
| Cross-validated R², soil profiles | 0.13 ± 0.12 | 0.08 ± 0.11 |
| Cross-validated R², studies | 0.03 ± 0.05 | 0.06 ± 0.09 |
| RMSE, random observations | 0.16 ± 0.00 | 0.95 ± 0.02 |
| RMSE, soil profiles | 0.21 ± 0.01 | 1.09 ± 0.07 |
| RMSE, studies | 0.22 ± 0.01 | 1.11 ± 0.05 |
| Number of soil profiles | 90 | 63 |
| MetaForest out-of-bag R² | 0.37 | 0.20 |

Table S6 Robustness of the main results. LOSO: maximum p-value obtained when each study is removed in turn.

| **Response** | **Test** | **Estimate** | **LOSO max p** | **Additional checks** | **Assessment** |
| --- | --- | --- | --- | --- | --- |
| Response ratio | Initial SOC stock (linear) | −1.8 percentage points per 10 Mg C ha⁻¹ [−2.7; −1.0]; p < 0.001 | 0.001 | ≥ 9 studies in each class of initial stock | Robust |
| Response ratio | Initial SOC stock × temperature | LRT = 11.5, df = 1, p < 0.001; slope −3.5% (28 °C) vs −1.5% (22 °C) per 10 Mg C ha⁻¹ | 0.010 | Accounting for agroforestry type: p < 0.001 | Robust |
| Response ratio | Temperature (spline) | LRT = 11.6, df = 3, p = 0.009 | 0.045 | Accounting for agroforestry type: p = 0.015; for type, initial SOC and precipitation: p = 0.19 | Not separable from confounders |
| Response ratio | Depth × precipitation | LRT = 10.9, df = 1, p < 0.001 | 0.034 | Deep observations (≥ 30 cm): 99 from 16 studies, 3–19 years | Exploratory |
| Response ratio | Systems ≤ 3 years vs older | 7.8% [−3.3; 20.1] vs 6.7% [1.6; 12.2]; interaction with design p = 0.75 | – | Accounting for initial SOC and temperature: p = 0.61 | No difference |
| Response ratio | Grey literature vs articles | LRT = 0.61, p = 0.43 | – | Grey literature: 93 observations from 13 studies | No difference |
| Storage rate | Initial SOC stock (linear) | −0.07 Mg C ha⁻¹ yr⁻¹ per 10 Mg C ha⁻¹ [−0.13; −0.01]; p = 0.025 | 0.10 | Spline: p < 0.001; three studies whose removal makes p > 0.05 | Significant but study-dependent |
| Storage rate | Temperature (spline) | LRT = 17.4, df = 3, p < 0.001 | 0.006 | Accounting for type, initial SOC and precipitation: p < 0.001; without sites > 28 °C: p = 0.014 | Robust, non-monotonic |
| Storage rate | Initial SOC stock × temperature | LRT = 0.97, p = 0.33 | – | – | Not supported |
| Storage rate | Depth × precipitation | LRT = 1.7, p = 0.19 | – | – | Not supported |
| Storage rate | Systems ≤ 3 years vs older | 0.59 [−0.06; 1.23] vs 0.24 [0.03; 0.45] Mg C ha⁻¹ yr⁻¹ | – | Accounting for initial SOC and temperature: p = 0.16 | Not significant |
| Storage rate | Grey literature vs articles | LRT = 1.93, p = 0.17 | – | Grey literature: 74 observations from 11 studies | No difference |

Table S7 Standardized XGBoost predictions by category (25 years after conversion for response ratios and 5 years for storage rates, other moderators held at their median or modal value).

| **Response** | **Moderator** | **Level** | **Prediction [95% interval]** | **Observations / studies** | **Maximum age observed (years)** | **Support** |
| --- | --- | --- | --- | --- | --- | --- |
| Response ratio (25 years) | Agroforestry type | Alley cropping/hedgerow | 1.14 [1.05; 1.22] | 43 / 8 | 31 | Yes |
| Response ratio (25 years) | Agroforestry type | Multistrata | 1.11 [1.02; 1.19] | 31 / 8 | 36 | Yes |
| Response ratio (25 years) | Agroforestry type | Other | 1.09 [1.01; 1.19] | 3 / 2 | 21 | No |
| Response ratio (25 years) | Agroforestry type | Shaded perennial crops | 1.17 [1.05; 1.27] | 71 / 13 | 20 | No |
| Response ratio (25 years) | Agroforestry type | Silvopasture | 1.15 [1.06; 1.28] | 170 / 29 | 70 | Yes |
| Response ratio (25 years) | Previous land use | Cropland | 1.17 [1.08; 1.27] | 75 / 13 | 37 | Yes |
| Response ratio (25 years) | Previous land use | Forest | 1.19 [1.08; 1.30] | 13 / 4 | 13 | No |
| Response ratio (25 years) | Previous land use | Grassland | 1.18 [1.03; 1.29] | 78 / 7 | 20 | No |
| Response ratio (25 years) | Previous land use | Unknown or mixed | 1.13 [1.04; 1.23] | 152 / 30 | 70 | Yes |
| Response ratio (25 years) | Tree species added | One species | 1.15 [1.06; 1.24] | 254 / 39 | 70 | Yes |
| Response ratio (25 years) | Tree species added | Two or more species | 1.15 [1.04; 1.33] | 64 / 17 | 40 | Yes |
| Storage rate (5 years) | Agroforestry type | Alley cropping/hedgerow | −0.09 [−0.57; 0.53] | 41 / 7 | 31 | Yes |
| Storage rate (5 years) | Agroforestry type | Other | −0.10 [−0.61; 0.59] | 11 / 4 | 36 | Yes |
| Storage rate (5 years) | Agroforestry type | Shaded perennial crops | 0.08 [−0.54; 0.63] | 60 / 10 | 20 | Yes |
| Storage rate (5 years) | Agroforestry type | Silvopasture | −0.03 [−0.50; 0.60] | 136 / 23 | 70 | Yes |
| Storage rate (5 years) | Previous land use | Cropland | 0.04 [−0.54; 0.63] | 67 / 12 | 37 | Yes |
| Storage rate (5 years) | Previous land use | Forest | −0.00 [−0.40; 0.69] | 13 / 5 | 13 | Yes |
| Storage rate (5 years) | Previous land use | Grassland | 0.06 [−0.41; 0.67] | 81 / 8 | 20 | Yes |
| Storage rate (5 years) | Previous land use | Unknown or mixed | −0.11 [−0.62; 0.48] | 87 / 18 | 70 | Yes |
| Storage rate (5 years) | Tree species added | One species | −0.03 [−0.54; 0.57] | 209 / 32 | 70 | Yes |
| Storage rate (5 years) | Tree species added | Two or more species | 0.02 [−0.49; 0.67] | 39 / 11 | 37 | Yes |

Table S8 Sensitivity of the storage-rate results to a 50% inflation of imputed standard deviations. Response-ratio results are unaffected because their sampling variances are computed from reported means, standard deviations and replicate numbers without imputation-specific scaling. Machine-learning results are identical in the two specifications because gradient-boosted trees do not use the sampling variances.

| **Quantity** | **Main analysis** | **Imputed SDs × 1.5** |
| --- | --- | --- |
| Overall storage rate (Mg C ha⁻¹ yr⁻¹) | 0.25 [0.05; 0.45] | 0.26 [0.06; 0.46] |
| Topsoil (0–30 cm) | 0.24 [0.01; 0.46] | 0.25 [0.01; 0.48] |
| I² (%) | 92.7 | 92.3 |
| Between-study share of heterogeneity (%) | 75.1 | 76.3 |
| Heterogeneity explained, biophysical (%) | 22.7 (LRT = 47.9, df = 13, p < 0.001) | 22.1 (LRT = 46.6, df = 13, p < 0.001) |
| Heterogeneity explained, system descriptors (%) | 0.0 (p = 0.77) | 0.0 (p = 0.80) |
| Initial soil carbon, linear | slope −0.068 [−0.128; −0.009], p = 0.025 | slope −0.083 [−0.145; −0.020], p = 0.010 |
| Initial soil carbon, spline | LRT = 18.4, df = 3, p < 0.001 | LRT = 18.0, df = 3, p < 0.001 |
| Initial soil carbon, leave-one-study-out | maximum p = 0.10 (3 studies) | maximum p = 0.049 (none) |
| Temperature, spline | LRT = 17.4, df = 3, p < 0.001 | LRT = 16.9, df = 3, p < 0.001 |
| Temperature given system type, initial stock and precipitation | LRT = 17.3, p < 0.001 | LRT = 17.0, p < 0.001 |
| Time since conversion, spline | LRT = 7.1, df = 3, p = 0.068 | LRT = 7.7, df = 3, p = 0.054 |
| Sampling depth, spline | LRT = 1.9, df = 3, p = 0.60 | LRT = 2.1, df = 3, p = 0.54 |
| Study design, mean response | LRT = 3.5, df = 2, p = 0.18 | LRT = 3.6, df = 2, p = 0.17 |
| Randomized versus before–after, standard deviation (%) | +98 | +139 |
| Change in residual standard deviation, 2 to 10 years (%) | −31 | −31 |
| Spanish versus English | LRT = 0.9, p = 0.33 | LRT = 1.0, p = 0.31 |
| Grey versus peer-reviewed | LRT = 1.9, p = 0.17 | LRT = 2.3, p = 0.13 |
| Egger-type test (p) | 0.54 | 0.67 |

Table S9 Articles present in the deposited database but not used in the analyses. The database is broader than this meta-analysis: it also holds comparisons between agroforestry systems and land-use-change experiments, which are outside its scope. The full reconciliation between the database and the two analysis datasets is given in outputs/data/database_vs_analysis_datasets.csv in the code repository.

| **Study** | **Rows in database** | **Rows with a tree-less control** | **Reason not used in the analyses** |
| --- | --- | --- | --- |
| 11 | 6 | 0 | No tree-less control; only comparisons between agroforestry systems or land-use-change experiments |
| 18 | 6 | 0 | No tree-less control; only comparisons between agroforestry systems or land-use-change experiments |
| 34 | 75 | 15 | Incubation study (ground tree litter incubated in jars): no field-based tree-less control |
| 35 | 27 | 0 | No tree-less control; only comparisons between agroforestry systems or land-use-change experiments |
| 38 | 6 | 0 | No tree-less control; only comparisons between agroforestry systems or land-use-change experiments |
| 62 | 6 | 0 | No tree-less control; only comparisons between agroforestry systems or land-use-change experiments |
| 64 | 2 | 0 | No tree-less control; only comparisons between agroforestry systems or land-use-change experiments |
| 72 | 18 | 0 | No tree-less control; only comparisons between agroforestry systems or land-use-change experiments |
| 83 | 28 | 0 | Control defined by distance from trees, not an independent tree-less plot |
| 86 | 2 | 0 | No tree-less control; only comparisons between agroforestry systems or land-use-change experiments |
| 88 | 2 | 2 | No comparison usable for an effect size; dispersion or replicate numbers not reported, or flagged as not usable at extraction |
| 96 | 3 | 0 | No tree-less control; only comparisons between agroforestry systems or land-use-change experiments |
| 107 | 18 | 0 | No tree-less control; only comparisons between agroforestry systems or land-use-change experiments |
| 131 | 1 | 0 | No tree-less control; only comparisons between agroforestry systems or land-use-change experiments |
| 138 | 12 | 12 | No comparison usable for an effect size; dispersion or replicate numbers not reported, or flagged as not usable at extraction |
| 157 | 3 | 0 | No tree-less control; only comparisons between agroforestry systems or land-use-change experiments |
| 172 | 9 | 0 | No tree-less control; only comparisons between agroforestry systems or land-use-change experiments |
| 174 | 1 | 1 | No comparison usable for an effect size; dispersion or replicate numbers not reported, or flagged as not usable at extraction |
| 1480 | 2 | 2 | No comparison usable for an effect size; dispersion or replicate numbers not reported, or flagged as not usable at extraction |

Table S10 Absolute-scale analysis and correction for regression to the mean. The absolute difference is SOC in the agroforestry plot minus SOC in the paired control (Mg C ha⁻¹), weighted by its sampling variance and fitted with the same three-level structure as the main models. λ is the share of the variance of the control stocks due to measurement error; the corrected slope is (β + λ) / (1 − λ) (Blomqvist 1977; Slessarev et al. 2023). Corrected values are point estimates: the uncertainty of λ is not propagated.

| **Quantity** | **Estimate** | **Test** | **λ** | **Corrected slope** |
| --- | --- | --- | --- | --- |
| Mean absolute difference (Mg C/ha) | 2.86 [0.28; 5.45] | p = 0.0301 | – | – |
| Absolute difference ~ initial SOC (linear) | −0.0170 [−0.0636; +0.0296] | LRT = 0.50, df = 1, p = 0.48 | – | – |
| Absolute difference ~ initial SOC (spline) | – | LRT = 5.49, df = 3, p = 0.139 | – | – |
| Absolute difference ~ initial SOC, given time and temperature | −0.0037 [−0.0498; +0.0425] | LRT = 0.02, df = 1, p = 0.877 | – | – |
| Absolute difference ~ initial SOC x temperature | – | LRT = 1.86, df = 1, p = 0.173 | – | – |
| Absolute difference ~ time since conversion (spline), given stock and temperature | +0.1956 Mg C/ha/yr (linear term) | LRT = 15.36, df = 3, p = 0.00154 | – | leave-one-study-out maximum p = 0.0346 |
| Slope on initial SOC: All observations | −0.0169 [−0.0637; +0.0299] (n = 322, 52 studies) | p = 0.48 | 0.1062 | +0.0999 |
| Slope on initial SOC: Control-impact designs | −0.1459 [−0.2257; −0.0661] (n = 177, 35 studies) | p = 0.000339 | 0.1270 | −0.0216 |
| Slope on initial SOC: Before-after designs | +0.0652 [−0.0027; +0.1332] (n = 39, 7 studies) | p = 0.0599 | 0.0140 | +0.0803 |
| Slope on initial SOC: Randomized designs | +0.0490 [−0.0508; +0.1489] (n = 106, 10 studies) | p = 0.336 | 0.0717 | +0.1300 |
| Slope on initial SOC: Surface layers (0-30 cm) | −0.0596 [−0.1206; +0.0015] (n = 167, 46 studies) | p = 0.0557 | 0.0892 | +0.0325 |
| Slope on initial SOC: At least 3 replicates | −0.0244 [−0.0756; +0.0268] (n = 236, 40 studies) | p = 0.35 | 0.0381 | +0.0142 |

Table S11 Relative importance of the moderators (% of total importance) when the machine-learning models are fitted to the absolute difference between agroforestry and control instead of the response ratio. Both responses were fitted to the same 322 observations from 52 studies, with the same predictors and hyperparameters. MetaForest weights each observation by the inverse of its sampling variance and groups observations by experiment; XGBoost is unweighted, and its values are means over 100 cluster bootstrap resamples, with the 17th and 83rd percentiles in brackets. Outliers were not removed before fitting, so the response-ratio columns are not directly comparable with the importance values of Fig. 3a. The sampling variance of an absolute difference varies far more among observations than that of a log response ratio, which is why the unweighted trees and the variance-weighted forest rank the initial stock differently.

| **Moderator** | **MetaForest, absolute difference** | **MetaForest, response ratio** | **XGBoost, absolute difference** | **XGBoost, response ratio** |
| --- | --- | --- | --- | --- |
| Temperature | 25.9 | 23.3 | 9.0 [5.1; 12.6] | 11.2 [6.7; 16.2] |
| Time since conversion | 21.5 | 14.7 | 14.9 [6.2; 24.4] | 12.3 [7.7; 16.9] |
| Precipitation | 13.7 | 14.8 | 13.0 [6.9; 18.4] | 14.7 [9.0; 20.0] |
| Main crop | 11.7 | 8.5 | 10.7 [1.0; 19.8] | 4.3 [1.6; 7.0] |
| Initial SOC stock | 6.8 | 18.1 | 34.4 [28.3; 41.4] | 40.3 [32.2; 49.2] |
| Sampling depth | 6.6 | 3.3 | 6.7 [3.5; 9.5] | 6.7 [4.4; 8.8] |
| Previous land use | 4.2 | 5.9 | 3.3 [1.3; 5.2] | 4.3 [1.7; 6.6] |
| Agroforestry type | 3.9 | 3.1 | 5.3 [2.5; 7.7] | 3.4 [1.9; 5.2] |
| Tree species added | 2.9 | 2.8 | 2.1 [0.5; 3.5] | 2.2 [0.9; 3.1] |
| Study design | 2.7 | 5.4 | 0.6 [0.1; 1.0] | 0.7 [0.2; 1.2] |
| Out-of-bag R² | 0.088 | 0.204 | – | – |

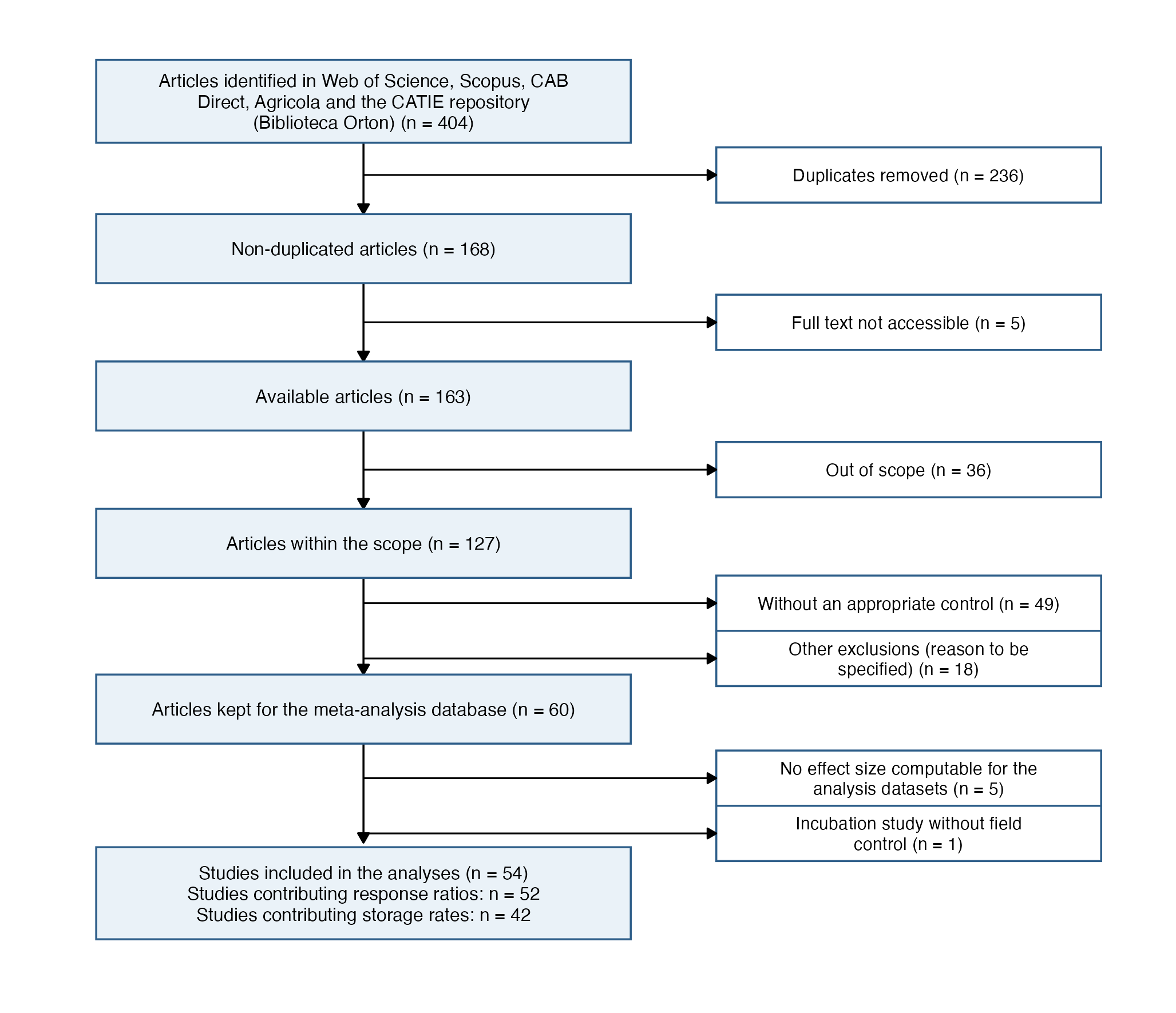

**Fig. S1** Flow diagram of the literature search, screening and inclusion of studies (PRISMA).

**Fig. S2** Experimental designs of the primary studies.

**Before–after designs** compare system states before and after the intervention. They account for temporal variability but lack control sites, making them susceptible to confounding factors unrelated to the intervention.

**Control–impact designs** compare impacted sites with non-impacted control sites at the same point in time. Because site allocation is typically non-random, differences between control and impact sites may partly reflect pre-existing spatial variation; this design is conceptually similar to space-for-time substitution.

**Randomized designs** randomly assign independent sampling units to treatment and control groups. Random allocation minimizes systematic differences between groups and reduces the need for pre-impact sampling.

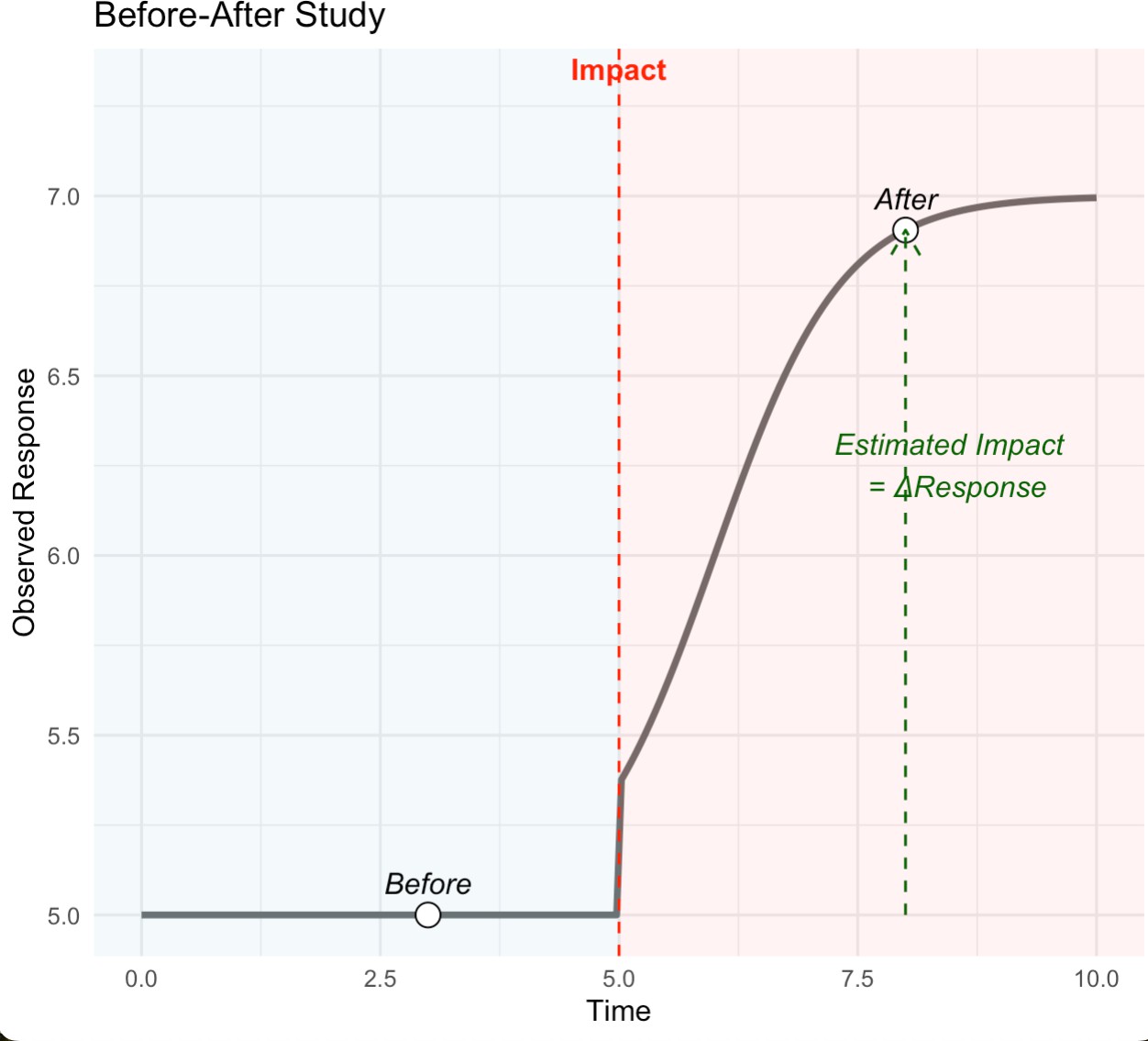

(a) Before–after design

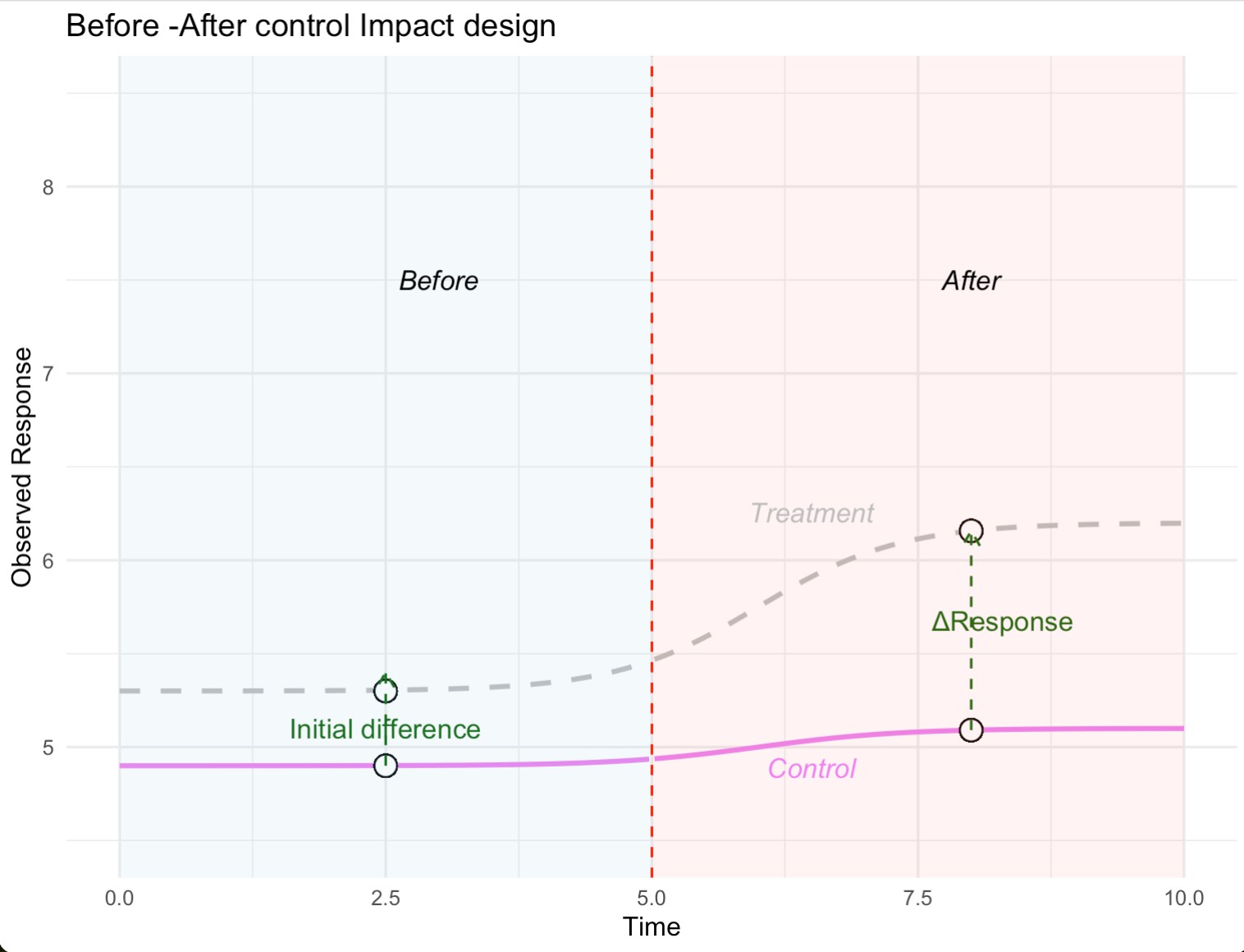

(b) Control–impact design

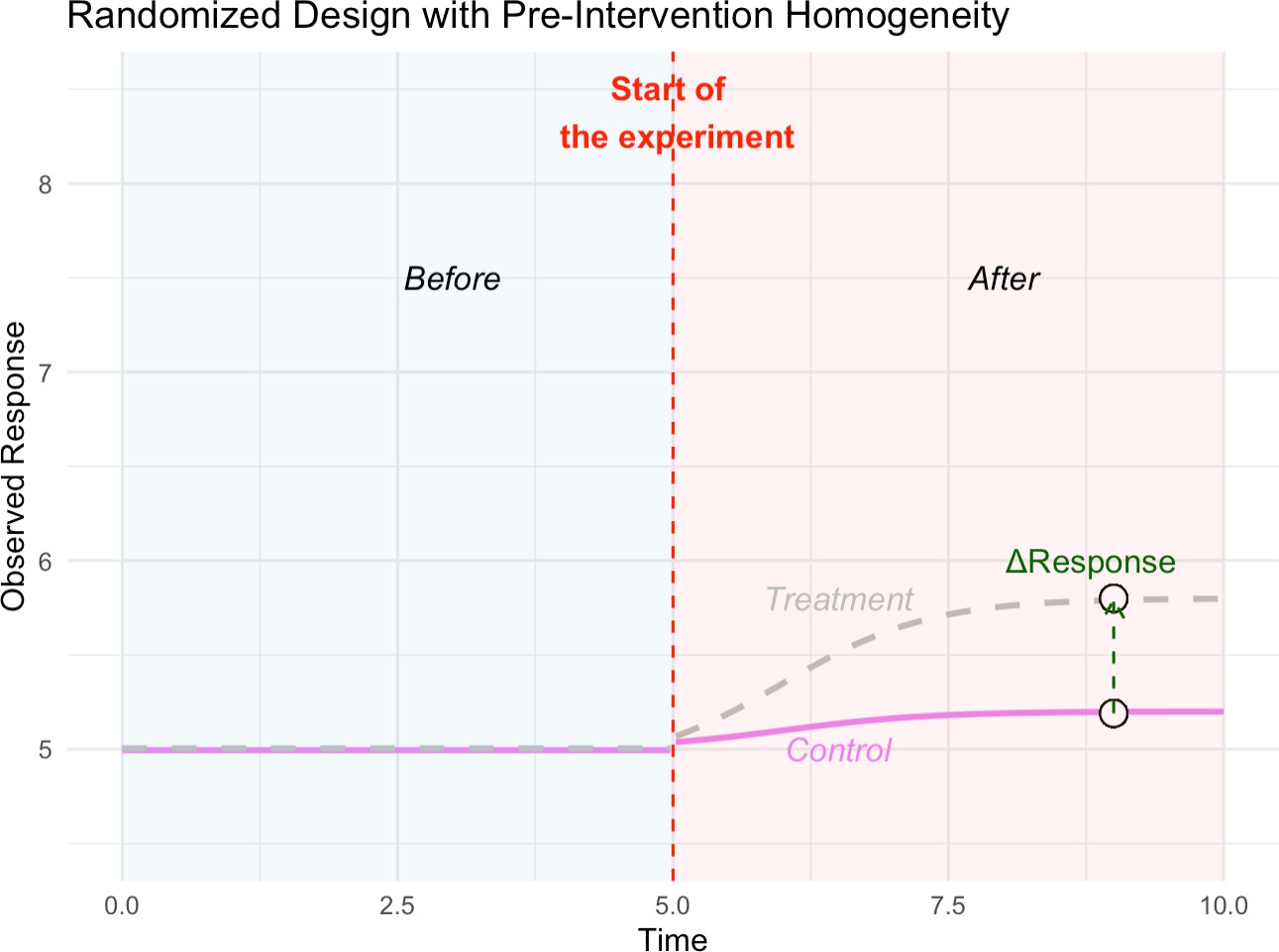

(c) Randomized design

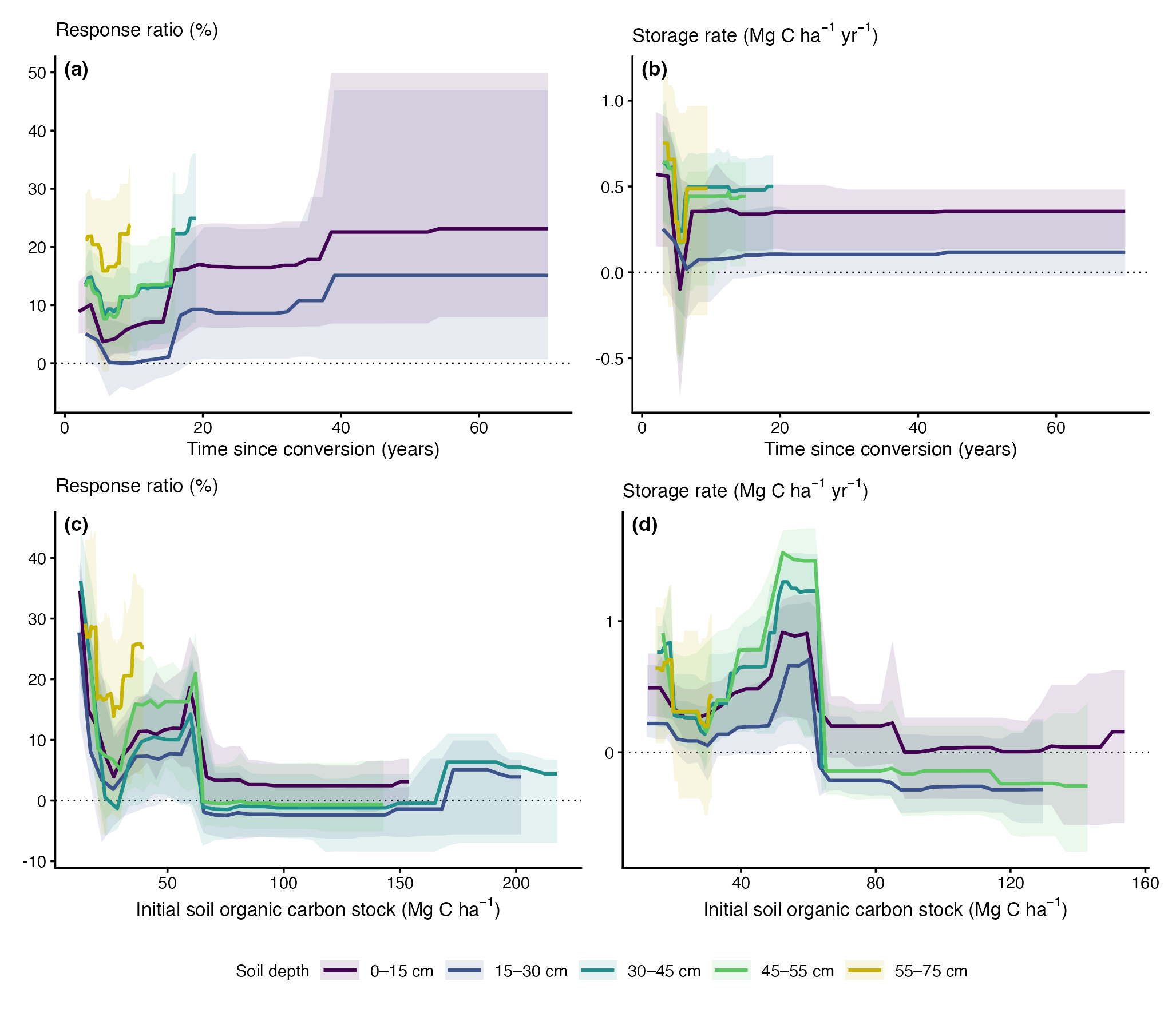

**Fig. S3** Partial dependence of XGBoost predictions on time since conversion (a, b) and initial soil organic carbon stock (c, d) for response ratios (a, c) and storage rates (b, d), by soil depth class. Curves are restricted to the range of each moderator observed within each depth class; bands show 95% intervals from 200 models refitted on bootstrap resamples of studies; depth classes are distinguished by line colour and listed in the legend. These model-based curves are descriptive; inference relies on the meta-regressions.

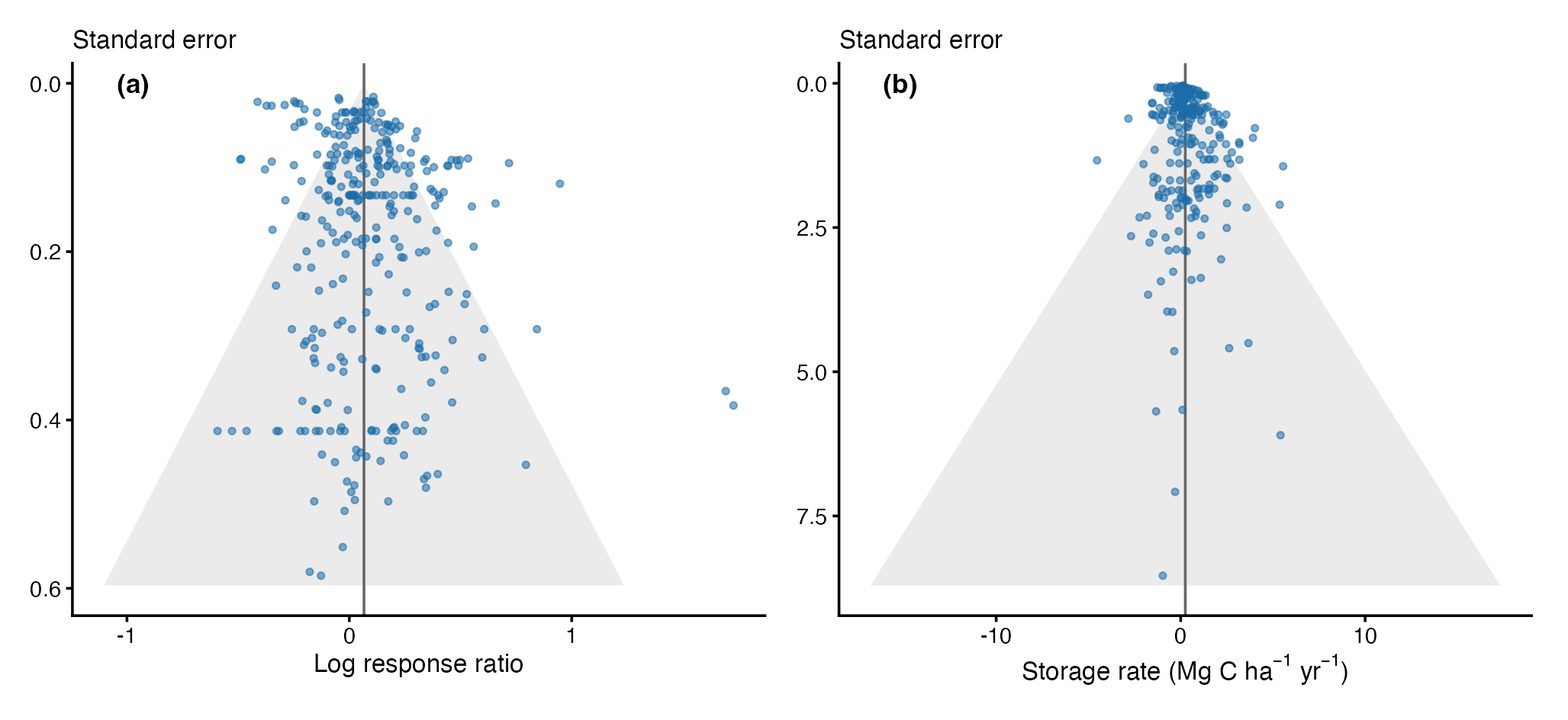

**Fig. S4** Funnel plots of (a) log response ratios and (b) storage rates against their standard errors. The vertical line shows the overall three-level estimate and dashed lines its pseudo 95% limits; symbols indicate study design. Multilevel Egger-type tests: p = 0.60 for response ratios and p = 0.40 for storage rates.

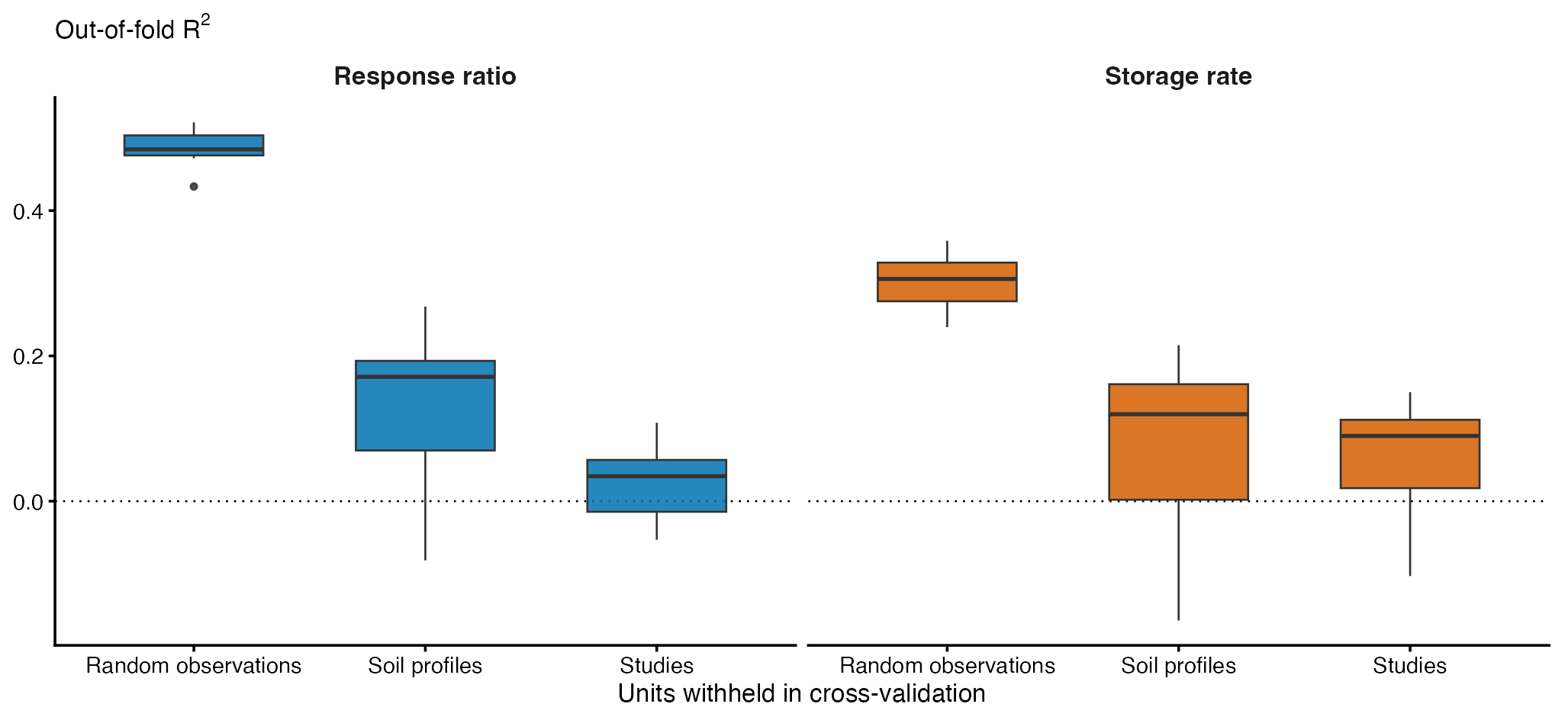

**Fig. S5** Out-of-fold R² of XGBoost models under three cross-validation schemes (ten repetitions of five-fold cross-validation): folds formed by random observations, by soil profiles (all depths of an agroforestry–control comparison kept together) and by studies.

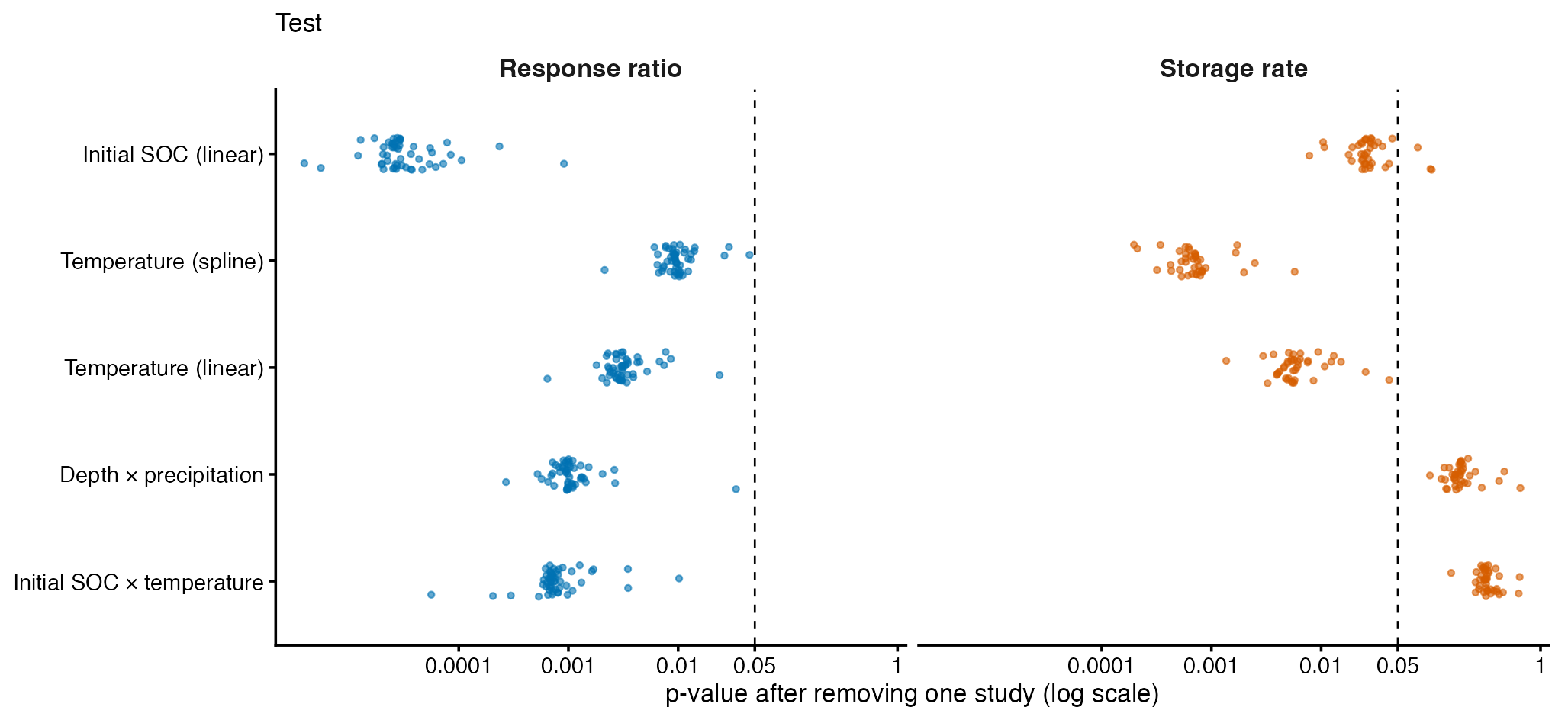

**Fig. S6** p-values of the robustness tests after removing each study in turn (leave-one-study-out), for response ratios and storage rates. The dashed line indicates p = 0.05; for storage rates, tests that were not significant with all studies are shown for completeness.

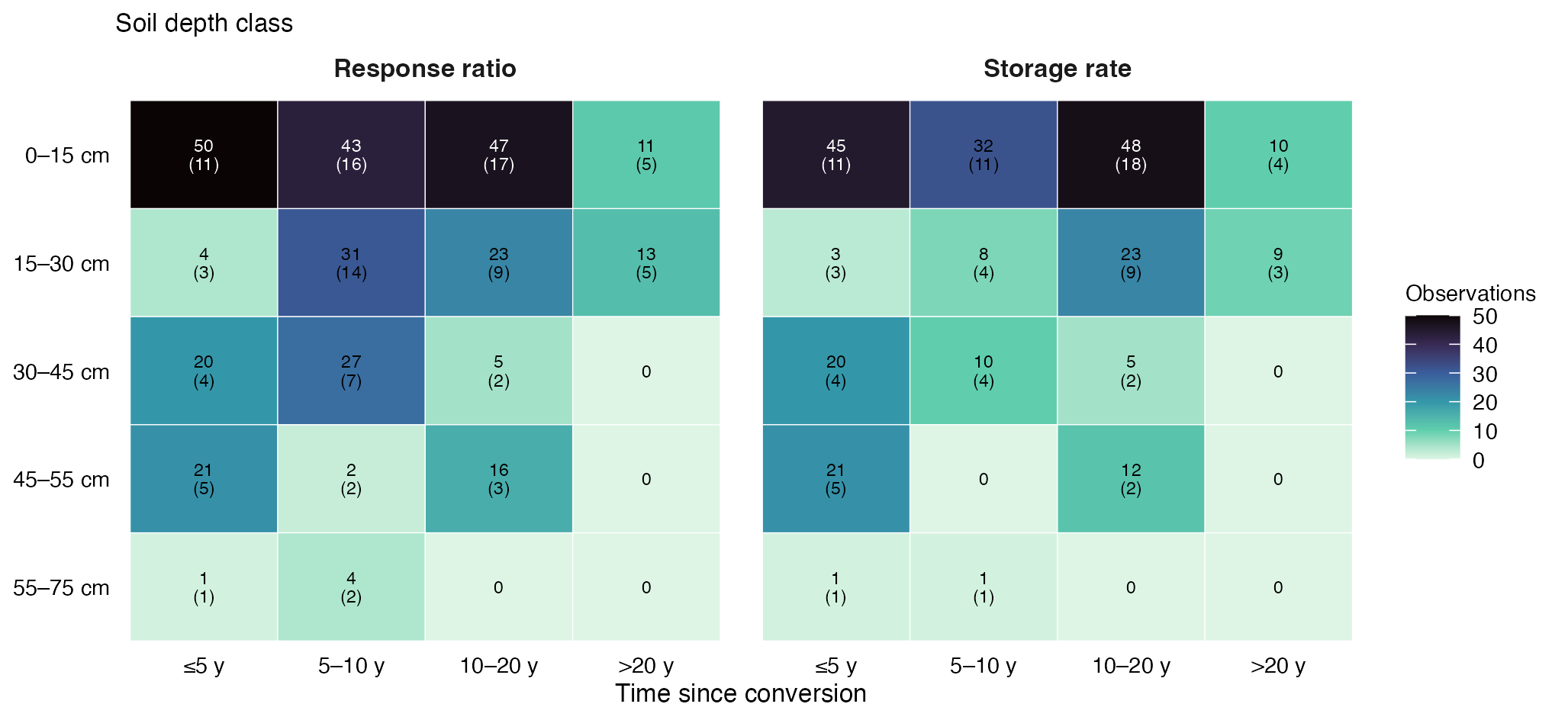

**Fig. S7** Number of observations (and of studies, in parentheses) by soil depth class and time since conversion.

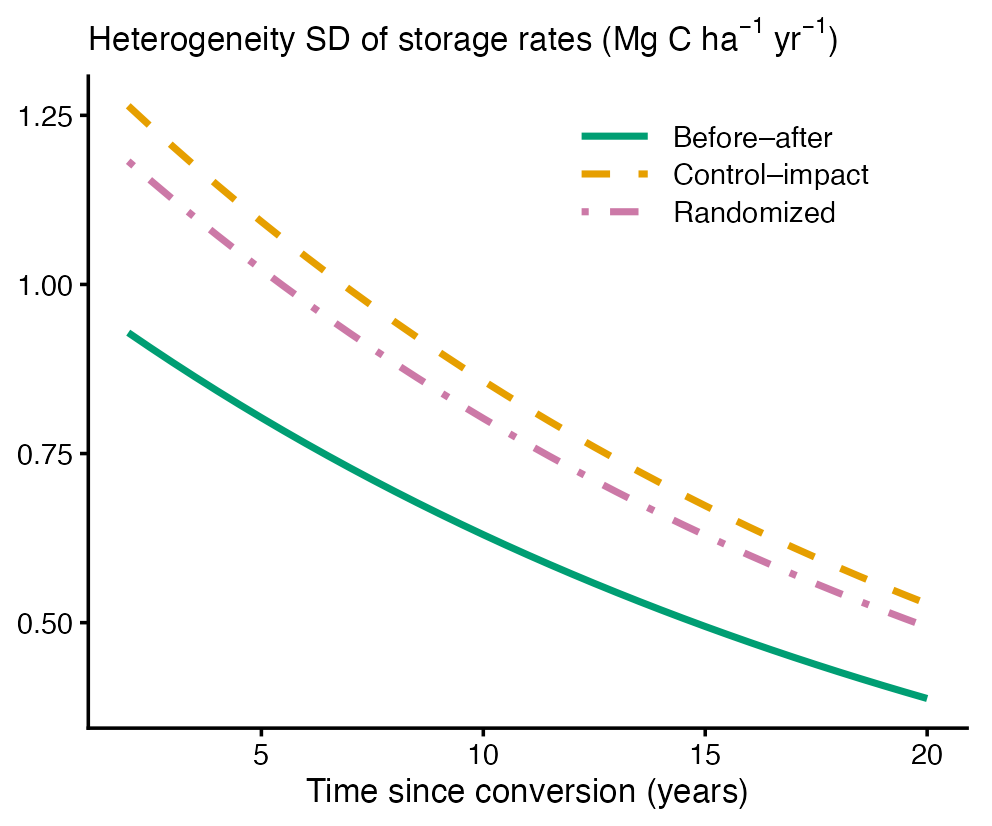

**Fig. S8** Standard deviation of true effects (residual heterogeneity) of storage rates as a function of time since conversion for before–after, control–impact and randomized designs, estimated with a location-scale model (Mg C ha⁻¹ yr⁻¹).
